# Dynamic modulation of auditory hair cell stereocilia membrane mechanics by the scrambling mechanotransduction complex

**DOI:** 10.1101/2024.09.27.615395

**Authors:** Shefin S. George, Anthony J. Ricci

## Abstract

Inner ear sensory hair cells convert mechanical deflection of their hair bundle into an electrical signal by gating novel mechanoelectrical transduction (MET) channels. Transmembrane channel-like proteins (TMCs) are a component of the MET machinery and part of a superfamily of molecules whose members are ion channels or lipid scramblases. As the stereocilia membrane modulates MET channels properties and the stereocilia membrane has different properties than the soma, we investigated the potential scramblase activity of the MET machinery using a novel viscosity sensor BODIPY 1c. Developmental, genetic, electrophysiological and pharmacological data demonstrate that both TMC1 and TMC2 are mechanosensitive scramblases as activity depends on MET channel being open. Blockage of scramblase activity unmasks a second process that increases effective membrane viscosity independent of MET machinery. Together our data reveal a dynamic regulation of stereocilia membrane that may underlie the speed and sensitivity of the MET process.

## Introduction

### Hair Cell Mechanotransduction

Hair cells are the mechanoreceptors of the auditory and vestibular systems used to convert sound stimulation and head positional information into electrical signals that are synaptically translated to the brain. The hair bundle is the mechanosensitive organelle that converts mechanical vibrations into electrical signals by a mechano-electrical transduction (MET) process^1^. The hair cell MET process is the fastest and most sensitive of mechanosensitive systems characterized to date. The hair bundle is comprised of an array of stereocilia, actin-filled microvilli, that increase in height in a staircase-like manner^2^. Cadherin based filamentous structures, termed tip links, connect shorter stereocilia to their neighboring taller stereocilia such that hair bundle deflection toward the tallest stereocilia pulls the tip link, exerting force at the stereocilia tip^3,4^. This force causes MET channels, located at the top of the shorter stereocilia, to open allowing for ion flux^5^. Multiple components of the MET channel complex interact with protocadherin 15 (PCDH 15), the lower component of the tip-link, supporting the hypothesis that unlike most mechanoreceptors, the channel molecules are directly tethered to an extracellular link^6–9^.

### MET channel Gating

A non-zero MET channel resting open probability suggests the tip-link is under tension, exerting a standing force onto the MET channels. How this force is generated is unknown. A gating spring theory suggests that force is transmitted via an elastic element directly to the gating molecule of the channel so that increasing tension in the spring opens the channel. The hair cell MET system is more complex as channel gating increases hair bundle compliance, modelled as an additional element, gating swing, that is in series with the spring^1,10^. This configuration accounts for a nonlinear force-displacement plot and allows for the estimation of the length of the gating swing, typically greater than 1 nm. This length is unlikely to be represented by a single molecule. The gating-spring model, along with a nonlinear force-displacement relationship has been identified in all hair cell types evaluated, even in drosophila hearing organs^11–15^. Along with the nonlinear force displacement plot, consistent between hair cell types is an adaptation process. Adaptation is defined as a shift in the current displacement plot that can open or close MET channels without directly altering the slope (sensitivity of the plot)^16–18^. At least two components of adaptation are described, a fast response with time constants that are sub millisecond that is independent of calcium entry and a slower process, typically in the 10s of milliseconds that is regulated via calcium^19–21^. The molecular underpinnings of this process(es) remain controversial.

### Membrane Modulation of Mechanotransduction

There is growing evidence for the modulation of hair cell MET by its surrounding lipid bilayer^20,22–1.25^. Initial data demonstrated the MET open probability sensitivity to divalent cations was regulated externally and followed an Eisenman series consistent with membrane charge screening^20^. GsMTX4, a known modulator of mechanically-gated channels^26,27^, further supported a role for the membrane bilayer shifting the hair cell current-displacement plot to the right and inhibiting resting open probability increases induced by depolarization and lowering external calcium^20^. The first direct evidence of membrane effects came from measurements of lipid diffusivity where stereocilia were 9x more diffusive than the soma and steady-state diffusivity was proportional to MET channel resting open probability^25^. Computational data implicated the bilayer as potentially contributing to adaptation, channel cooperativity and gating compliance^17,22,24,28,29^. Together these data suggest the stereocilia membrane properties are not simply passive and that they directly impact MET function.

### Molecules of MET complex

Multiple components of the MET channel complex have been identified^2,30^. Central to these are the transmembrane channel-like 1 and 2 proteins (TMC1 and TMC2), TMIE, LHFPL5 as well as CIB2^31–38^. TMIE is required for both trafficking molecules within the complex and is a critical component of the MET channel^32,39–41^. LHFPL5 may be a part of the force translation pathway as it interacts with both TMC molecules and PCDH15^7,40,42^. TMC molecules are putative pore forming subunits that are part of a broader family of molecules including the TMEM and OSC molecules which can be ion channels or lipid transporters, specifically membrane scramblases^43^. Recent evidence suggests that TMCs may have scramblase activity and posit this activity may be important for maintaining membrane homeostasis^44^.

We present data that the MET channel complex can directly regulate membrane mechanical properties through scramblase activity. We further demonstrate an active process increasing stereocilia viscosity that works in opposition to the scramblase activity to modulate stereocilia effective viscosity. Our data provides a new foundation upon which to investigate channel gating and processes related to adaptation, gating springs and even cochlear amplification.

## Results

### Stereocilia membrane is less viscous compared to supporting cells

To measure the cochlear cell’s membrane properties, we used a viscosity-sensitive “molecular rotor” BODIPY 1c (Fig. 1A) coupled with fluorescence lifetime imaging (FLIM). Molecular rotors are small synthetic fluorophores that exhibit viscosity-dependent fluorescence quantum yield and fluorescence lifetime i.e., the average time a fluorophore remains in the excited state^45–48^. Rotation or intramolecular twisting of the molecular rotor leads to non-radiative decay from the excited state back to the ground state. In an ordered or more viscous membrane, the non-radiative decay pathway is restricted, leading to an increase in the fluorescence intensity and fluorescence lifetime. In less viscous membranes, fluorescence intensity and lifetime decreases (Fig. 1B). To obtain concentration-independent measurements of viscosity, fluorescence lifetime of BODIPY 1c was used. More details on lifetime measurements, validation and calibration of BODIPY 1c and determination of optimal incubation concentration of BODIPY 1c in cochlear hair bundles are presented in the Methods and in Supplementary data, Fig.S1-2. Fig. 1B schematizes more viscous (red) and less viscous (green) membranes with corresponding lifetime curves that were obtained from liposomes created with sphingomyelin/cholesterol (viscous) or pure DOPC (less viscous). Data are presented as lifetime throughout but reflect effective viscosity. It is not possible to calibrate the sensor in the target system where uncontrolled variables like membrane proteins and cytoskeletal connections could alter the absolute value of the measured viscosity; however, our control data suggest the sensor reports the effective membrane viscosity. Fig. 1C presents example FLIM images from live P10 rat mid-apical inner hair cell bundles stained with BODIPY 1c at the locations indicated in the schematic, ostensibly the tops of the first row and second row stereocilia. The donut-like fluorescent rings attest to the membrane localization of the BODIPY 1c.

**Figure 1:**
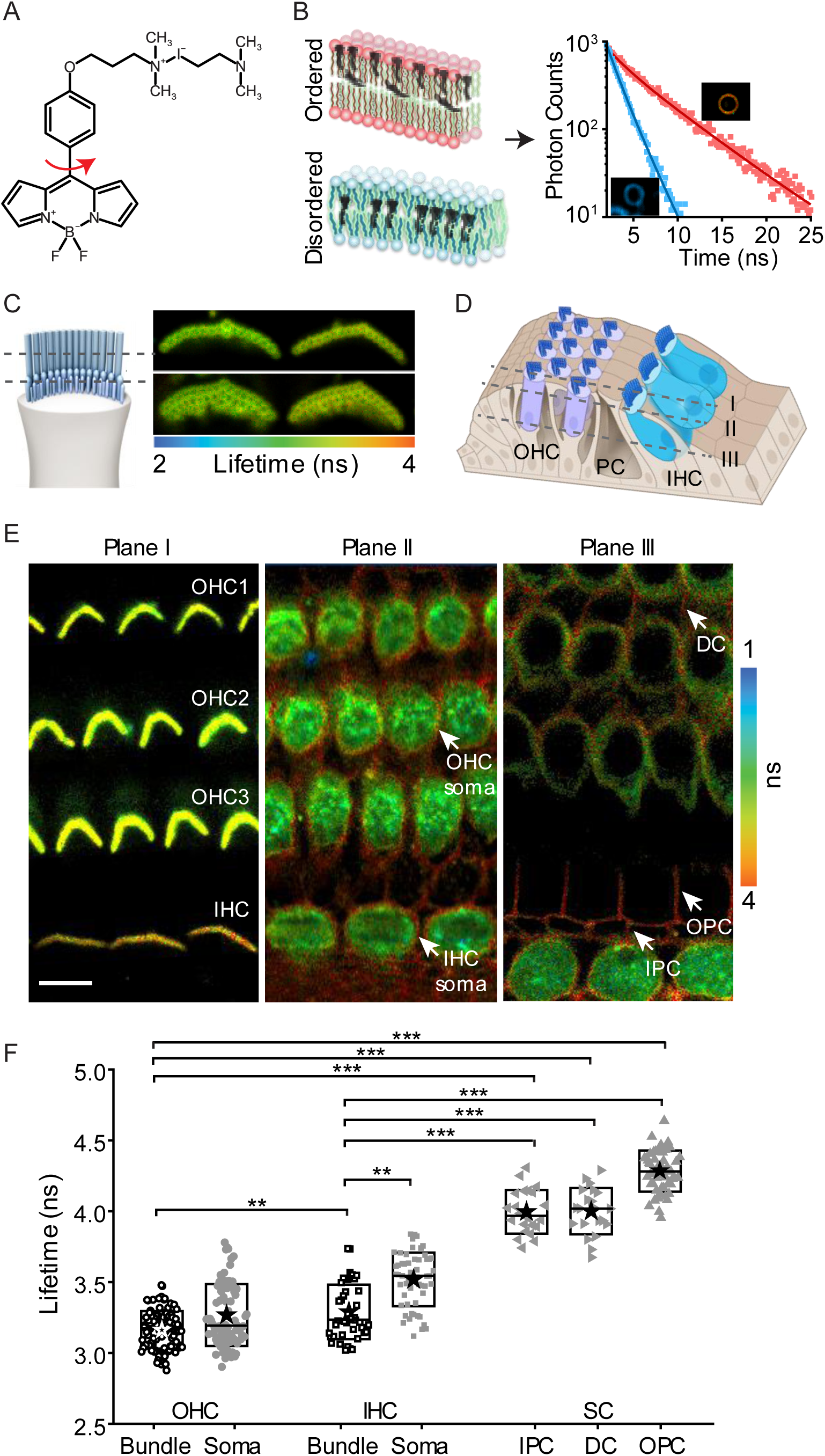
Stereocilia membrane is less viscous compared to pillar cells. A) Molecular structure of BODIPY 1c. B) Schematic showing the relative orientation of BODIPY 1c in an ordered and disordered bilayer (adapted from ^46^), example decay curves of BODIPY 1c from bilayer with low viscosity (blue symbols) and high viscosity (red symbols) and the corresponding exponential fits (solid lines) and example lifetime images generated with each pixel represented by the time constant (τ, mean lifetime) derived from fitting the decay curve from each pixel with a double exponential decay model. C) FLIM images of the inner hair cell (IHC) bundles of P10 rat mid-apical cochlear turn stained with BODIPY 1c with laser focused on the tops of the first and second row stereocilia with the schematic on the left showing the approximate location of the focal planes. D, E) Cross-sectional FLIM images of P10 rat mid-apical turn stained with BODIPY 1c in XY planes to present sensory cells (inner hair cells (IHCs) and outer hair cells (OHCs)) and supporting cells (inner pillar cells (IPCs), outer pillar cells (OPCs) and deiter cells (DCs)). D illustrates the approximate position of the focal plane of the FLIM images shown in E with grey dashed lines. F) Quantification of mean lifetime from different cochlear structures in P10 rat mid-apical turn. Boxes represent the SD, and the star symbol indicates the mean. Each data point corresponds to a hair bundle or a cell (for soma). \*\**p* < 0.01, \*\*\**p* < 0.001. Scale bar = 10 μm.

Previously, two-photon Fluorescent Recovery after Photobleaching (FRAP) was used to monitor membrane diffusivity, demonstrating that the stereocilia membrane is highly diffusive compared to the soma^25^. To investigate the viscosity differences between different structures of the organ of Corti, especially between the hair bundles and soma and between the sensory cells and the supporting cells, FLIM z-stacks of the organ of Corti were acquired. Fig. 1E shows cross- sectional FLIM images of P10 rat mid-apical turn stained with BODIPY 1c in XY planes to present sensory cells (inner hair cells (IHCs) and outer hair cells (OHCs)) and supporting cells (inner pillar cells (IPCs), outer pillar cells (OPCs) and deiter cells (DCs)). Fig. 1D illustrates the approximate position of the focal plane of the FLIM images shown in Fig. 1E organ of Corti with grey dashed lines. The fluorescence decay of BODIPY 1c in most cochlear structures, including the hair bundles and soma of HCs and supporting cells, at 20°C was best fitted with a biexponential model indicating that the rotor is probing the heterogeneities in the plasma membrane^45,49^. We confirmed the absence of dye aggregation, which could also render a biexponential decay^49^, as the two spectral windows for monomers and aggregates were identical at the dye concentration and incubation time used (see Supplementary data and Fig. S2A for more details). Using the time constants extracted from the biexponential decay, we report the intensity-weighted mean lifetime to represent the average lifetime. The measured lifetime of IHC bundles were significantly higher than that of the OHC bundles (*t*-test, *p<0.01*), and the sensory cells (IHCs and OHCs) bundles and soma were significantly lower than that of the supporting cells – IPCs, OPCs and DCs PCs (*t*-test, *p<0.001*) (Fig. 1F). For the IHCs, the bundles had significantly lower lifetime than that of the soma (*t*-test, *p<0.01*) (Fig. 1F).

For both IHC and OHCs, it was difficult to isolate the plasma membrane because the soma was filled with BODIPY 1c, likely underestimating the lifetime of hair cell soma plasma membrane.

### Membrane viscosity is significantly high in mice lacking MET channel complex

We first explored whether the lack of MET channel complex in the hair bundle would affect the stereocilia membrane effective viscosity. For this purpose, we captured FLIM images of organ of Corti from *Tmc1* and *Tmc2* double knockout mice (*Tmc1*^−/−^; *Tmc2*^−/−^), which lack functional MET channels^50^. TMC1 and TMC2 are implicated as pore-forming subunits of the MET channel^35,37,51–53^. In the *Tmc1^-/-^ ; Tmc2^-/-^* mice, the hair bundles in both the OHCs and the IHCs showed slower decay curves (Fig. 2B) and significantly higher lifetimes (Fig. 2A, C; *t*-test, *p<0.001*) compared to littermate controls, *Tmc1^-/+^* ;*Tmc2^-/+^*. The prominence of the fluorescence from the supporting cells in the double KOs is a consequence of the longer imaging time required to collect sufficient photons to monitor lifetime in the higher effective viscous conditions (as less dye is taken up). Also, the longer imaging times in the control mice would have saturated the hair cell cytoplasm signal contributing to the differences in images. No significant difference was observed between the PC soma of the *Tmc1^-/-^; Tmc2^-/-^* and the *Tmc1^-/+^; Tmc2^-/+^* mice suggesting that the effect on the membrane lifetime is specific to the HCs (Fig. 2C). A higher lifetime, indicative of a higher effective viscosity, is consistent with the loss of scramblase activity. As scramblases reduce membrane order, creating membrane heterogeneity, a reduction in effective viscosity is predicted. This contrasts with active transporters like flippases and flippases, where membrane order is increased, and thus effective viscosity would increase.

**Figure 2:**
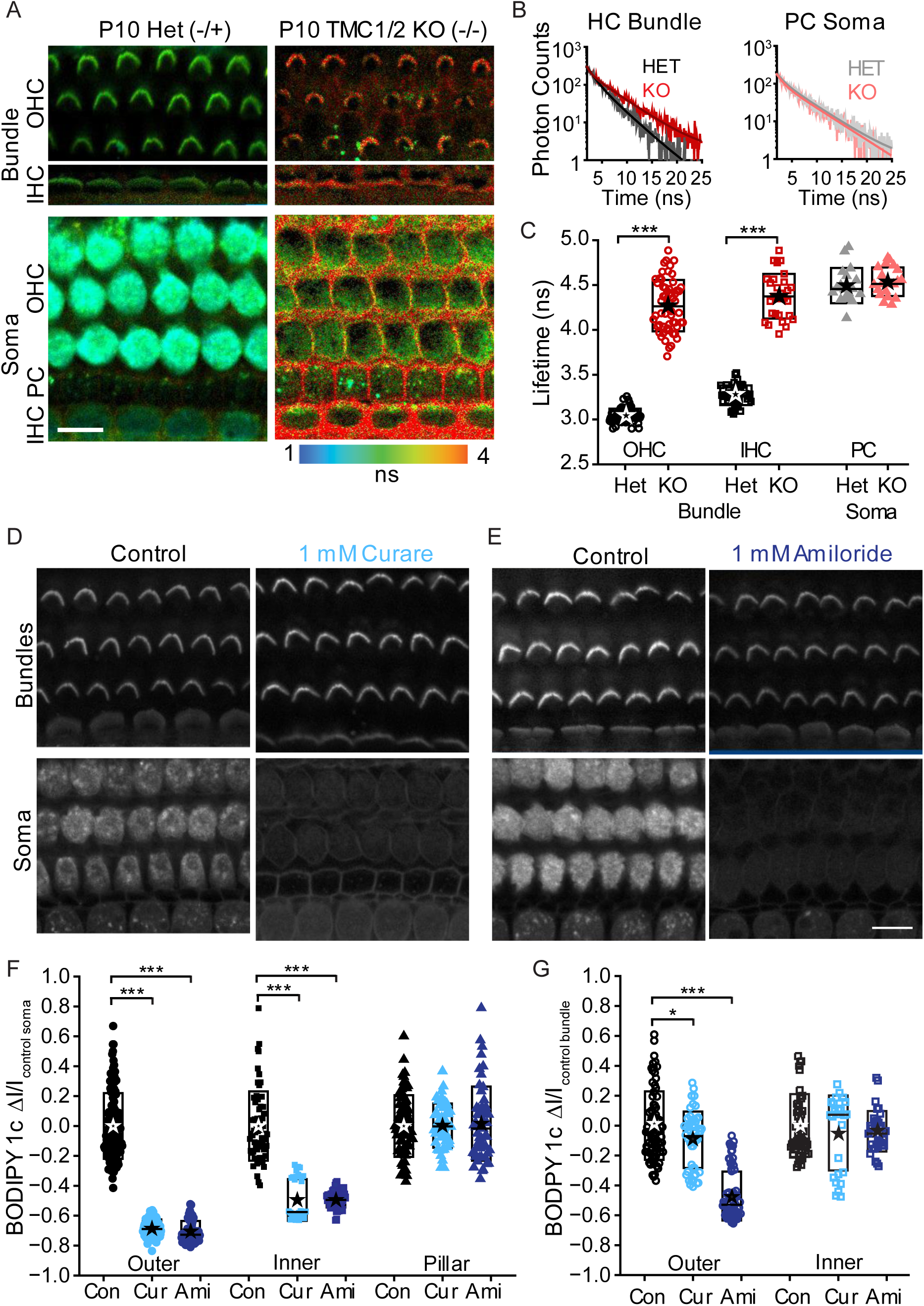
Membrane viscosity is significantly high in mice lacking MET channel. A) FLIM images of P10 *Tmc1*^−/−^;*Tmc2*^−/−^ and *Tmc1*^−/^ ;*Tmc2*^−/^ HC bundles (top rows) and soma (bottom row). B) Example fluorescence decay curves comparing the HC bundles and PC soma of double knockout mice and the littermate controls. C) Quantification of the mean lifetime from *Tmc1*^−/−^;*Tmc2*^−/−^ and littermate controls. D, E) Intensity images of BODIPY 1c in P10 rat mid-apical turn of control, 1mM curare and 1mM amiloride treated organ of Corti with focus plane at the hair bundles (top row) and soma (bottom panel). F, G) Quantification of BODIPY 1c intensity in the F) soma of the hair cells and G) the hair bundles. Boxes in C, F and G represent the SD, and the star symbol indicates the mean. Each data point corresponds to a hair bundle or a cell (for soma). \**p* < 0.05, \*\**p* < 0.01, \*\*\**p* < 0.001. Scale bar = 10 μm.

Given that BODIPY 1c was found in the cytoplasmic membranes of HCs in controls and not supporting cells or mutant HCs without MET channels, we examined whether the dye could enter the HCs through MET channels by monitoring fluorescence intensity in the soma treated with MET channel blockers, curare and amiloride^54–58^. Incubation with 1 mM curare or amiloride resulted in a significant reduction of intensity in the soma of HCs and not PCs (Fig. 2D-F, *t*-test, *p<0.001*), indicating that BODIPY 1c enters hair cells at least in part through the MET channel; a useful property as it allows hair cells with open MET channels to be identified. The cytoplasmic BODIPY 1c fluorescence also correlated with the cytoplasmic signal of FM 4-64, (Fig. S3A, B, *r2*=0.79) a fluorescent dye previously shown to enter hair cells via MET channels^41,59,60^. Thus, the HC soma membrane lifetime measurements reported in Fig. 1F are contaminated by the presence of BODIPY 1c in the cytoplasm which could underestimate the lifetime of the HC soma. For this reason, we did not pursue HC soma comparisons further. The images focused on the soma of the HCs (Fig. 2A; lower panels) showed the absence of BODIPY 1c in the cytoplasm of the double knockout mice compared to littermate controls, confirming that the dye did not enter the HC soma without the MET channel complex.

### Stereocilia membrane viscosity decreases during development

Cochlear hair bundles undergo structural and functional maturation over several days during development^61^. Along with the changes in stereocilia height, width and number of rows per hair bundle, mechanosensitivity matures over several days during development with a spatial and temporal progression of both functional and structural maturation of the hair bundle^61^. The expression pattern of essential MET components such as TMC1 and TMC2 vary during development, specifically within the cochlea, with TMC2 being expressed during early post-natal days but replaced over time by TMC1^31,38,50,62,63^. Therefore, we next explored whether membrane lifetime was impacted by MET using the natural temporal and spatial differences in MET maturation as a tool. Fig. 3 summarizes the results showing that stereocilia membrane lifetime co-varies with MET maturation. Three locations along the cochlea, apex, middle and base, were monitored from post-natal day 1 to 10 (P1-P10) rats. FLIM images in fig. 3A show that hair bundles that are not mechanically sensitive had higher lifetimes (indicative of higher viscosities). At P1, OHCs at apex and middle regions do not have MET currents and their hair bundles have higher lifetimes. The lack of MET currents is also illustrated by the absence of BODIPY1c in the hair cell soma. OHCs at the P1 base appear as a mosaic pattern where hair bundles span a broader range of lifetimes. Unexpectedly, IHC hair bundles had lower lifetimes in all regions with the apex having a mosaic pattern. Also, IHC soma were more filled at each position suggesting more functional MET in IHCs. Images taken at P3 show lower lifetimes in the base with the mosaic pattern having moved to the mid region and the apex remaining with higher lifetimes. By P7, hair bundles from all regions had lower lifetimes and soma were filled. These data are consistent with the hypothesis that membrane viscosity lowers as MET currents mature. Further support comes from the strong correlation between hair bundle lifetime and soma uptake of the sensor (Fig. 3D) measured from the mid-apex region. To more directly demonstrate that hair bundle lifetime correlated with MET maturation, we first captured FLIM images and then used whole-cell recordings to voltage clamp hair cells and measure maximal MET currents using a fluid jet stimulator^64^. P3 mid-apex was selected as there was a broad range of lifetimes for the hair bundles from that position. The plot in fig. 3F suggests that lifetimes above 5 ns are from hair bundles that have no MET currents. Figure 3E summarizes the developmental changes in bundle lifetime for the mid-apical turn. Both IHC and OHC bundles show lifetimes that reduce over time, where IHC bundles’ lifetime reduces at least 1 day earlier than OHC bundles. During this same time course, no changes in lifetime were found in PCs, again showing hair cell selectivity.

**Figure 3:**
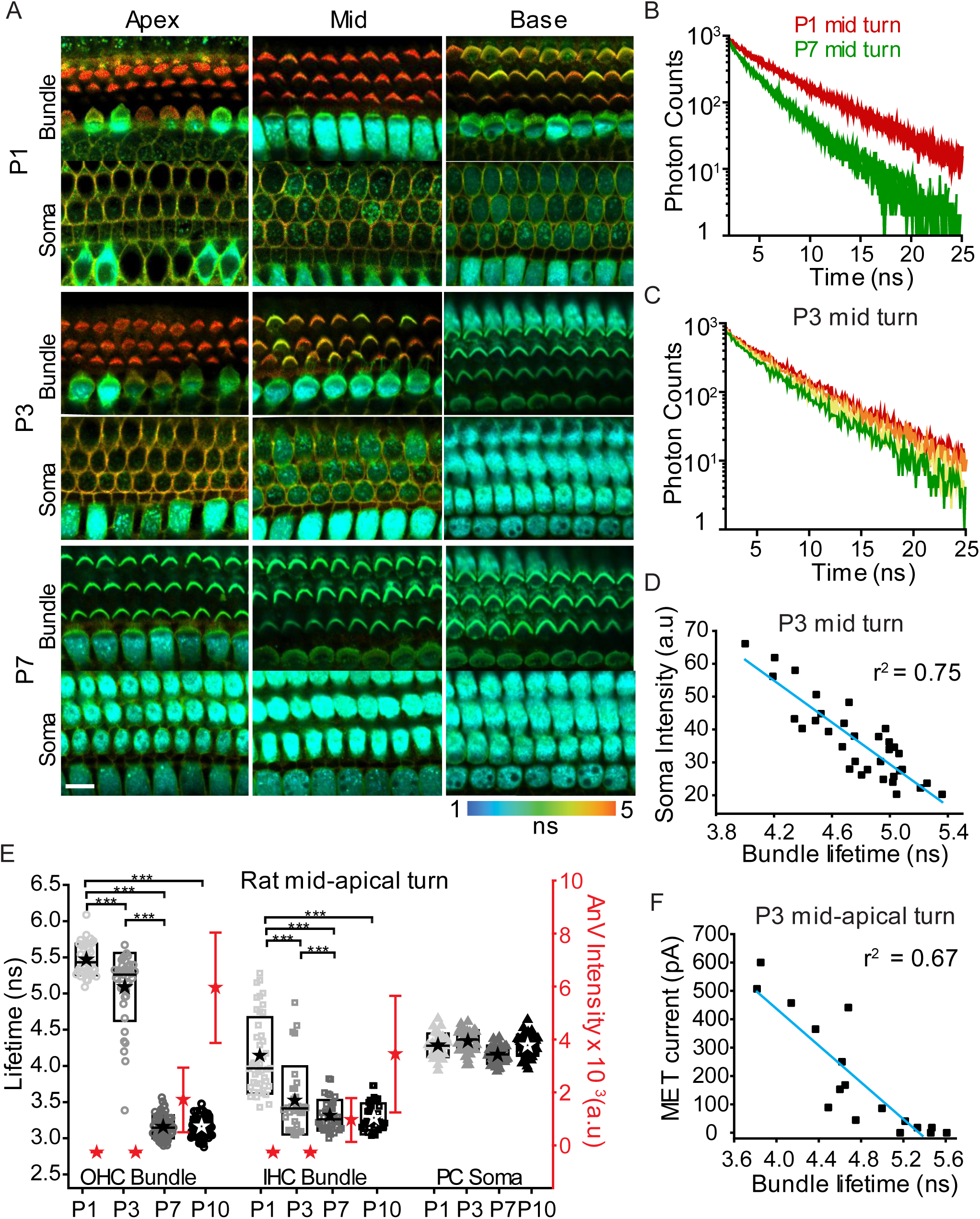
Stereocilia membrane viscosity decreases during development and correlates with the onset of MET. A) FLIM images of hair bundles and soma of developing rat organ of Corti stained with BODIPY 1c from three positions (apex, middle, base) along the cochlea at ages from postnatal day 1 (P1) to P7. B) Fluorescence decay from P1 (red traces) and P7 (green traces) hair bundles from middle turn. C) Fluorescence decay from P3 middle turn hair bundles of mosaic lifetimes with red, yellow and green traces from bundles with high, intermediate and low lifetimes respectively. D) Plot showing the correlation between the fluorescence intensity of BODIPY 1c in the soma correlates with the bundle lifetime. E) Time course of the change in lifetime (left axis) and PS externalization (right axis) for mid-apical OHC bundles (open circle), IHC bundles (open square) and PC soma (closed triangle). F) Plot showing the correlation between the bundle lifetime and the MET current measured from the corresponding cell. Boxes in (E) represent the SD, and the star symbol indicates the mean and the line indicates the median. Each data point corresponds to a hair bundle or a cell (for soma). \*\*\**p* < 0.001. Scale bar = 10 μm.

To test more directly if expression of TMC1 protein correlates with hair bundle lifetime, we tracked TMC1 endogenous protein using *Tmc1^HA/HA^* knock-in mice (Fig. S4). We first captured the FLIM images and then performed anti-HA immunofluorescence staining on the same sample of mid-apical OHC and imaged hair bundles with high-resolution confocal Lightning imaging to detect TMC1 at P4 in littermates and used non-HA animals as negative controls (Fig. S4A, B). In the control animals (*Tmc1^HA/HA^*), TMC1-HA was detected at low levels in the hair bundle and formed puncta along the stereocilia (Fig. S4A, C). The plot in Fig. S4D shows a strong correlation (r^2^ = 0.83) between the normalized sum of the HA fluorescence intensity along the hair bundle and the bundle lifetime.

We hypothesized that the changes in lifetime during development, which reflect changes in membrane viscosity, are due to the lipid scramblase activity associated with the MET machinery. As externalization of phosphatidylserine (PS), which is usually restricted to the inner leaflet, is often used as a marker for scramblase activity^65^, we used the PS-specific binding protein Annexin V (AnV) labeling to detect PS externalization. Fig. S5A shows AnV labeling for the mid-apical region between P1-P10. An increase in labeling occurs for both IHC and OHC bundles, summarized in Fig. S5B. As an aside, labeling at P7 clearly targeted remaining Kinocilia that are destined to be removed, this labeling is unlikely driven by TMCs. When correlating AnV labeling with the change in lifetime, most of the lifetime changes occur prior to the increase in AnV labeling (Fig. S5C). Fig. 3E exemplifies the difference in timing by including both lifetime (left axis) and AnV labeling (right axis). In considering scramblase activity, PS is not the only lipid transported and potentially other lipids such as phosphatidylethanolamine (PE) are scrambled earlier than PS^66–68^. However, given that there are many changes happening during development, it is difficult to directly interpret the difference in timing.

### Membrane viscosity is significantly high in mice lacking TMC1

We next wanted to determine if the TMC molecules were directly involved in the changes in membrane viscosity. We first chose to use the TMC1 KO mouse as the later onset of TMC1 allows for normal development to happen, including functional MET channels, thus allowing for the isolation of the TMC1 effect. Fig. 4 shows that, at both the apical and basal cochlea turns, hair bundle lifetime appears normal before P4 where TMC2 is expressed. However, lifetimes increase in the *Tmc1^-/-^* mice after P4 implicating TMCs as part of the membrane regulatory pathway. Fig. 4A to C show FLIM images and measured lifetimes for the mid-apex region, while Fig.4D to F show similar measurements for the basal regions. In the mid-apical turn, MET onset is late, as reflected in the membrane lifetime. Lifetime starts high and rapidly reduces between P1 and P4 (Fig. 4C). The littermate *Tmc1^-/+^* mice maintain lower lifetimes after P4 whereas the *Tmc1^-/-^* mice show an increase in lifetime for both IHCs and OHCs. Similarly, in the basal region where MET onset happens at or before P1, lifetimes are low and are maintained low for littermate controls; however, the *Tmc1^-/-^* have an increasing lifetime (Fig. 4F). These data show a direct link between TMC expression and membrane lifetime (effective viscosity). However, it was surprising that lifetimes in the *Tmc1^-/-^* did not go back to pre-MET onset levels (Fig. 4C, F).

**Figure 4:**
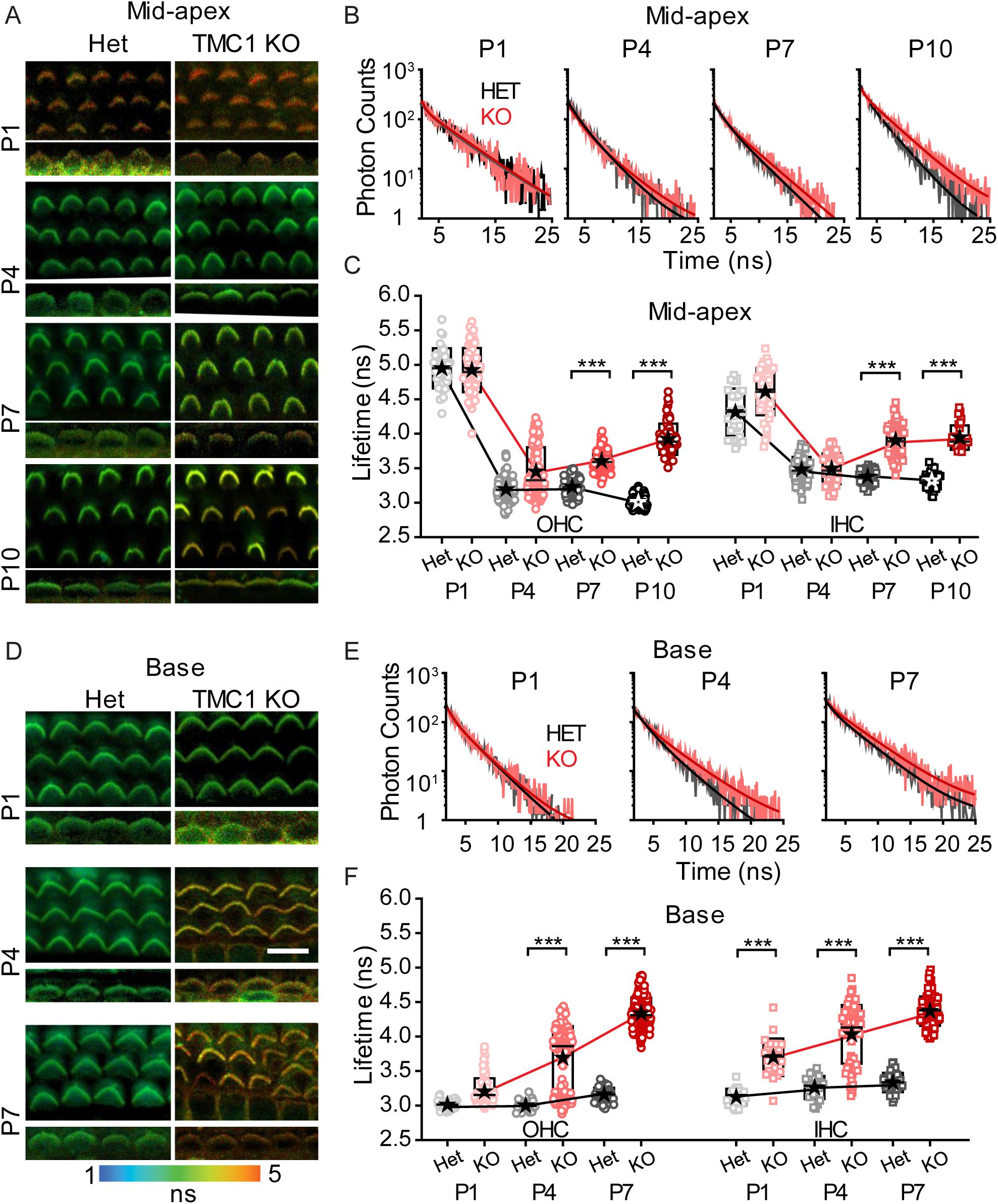
TMC1 and TMC2 are required for the reduction of membrane viscosity during development. FLIM images of OHC and IHC bundles from A) mid-apical turn of P1, P4, P7 and P10 and D) basal turn of P1, P4 and P7 mice cochlea from *Tmc1*^−/−^ and *Tmc1*^−/^ . Example decay data and corresponding fitted curve comparing the *Tmc1*^−/−^ and controls for mid-apical turn (B) and basal turn (E). The measured hair bundle lifetime for both controls and *Tmc1*^−/−^ from C) mid-apical turn and F) basal turn. Boxes in (C) and (F) represent the SD, and the star symbol indicates the mean and the line indicates the median. Each data point corresponds to a hair bundle. \*\*\**p* < 0.001. Scale bar = 10 μm.

Assessment of hair cell soma dye uptake suggests MET current is still present at a lower level in the *Tmc1^-/-^*. Fig. S6 summarizes this data from P10 *Tmc1^-/-^* and littermate controls (*Tmc1^-/+^*) for IHC and OHC bundle and PC soma. The summary plot of Fig. S6C shows the increase in membrane lifetime for both IHC and OHC bundles but not for PCs. Inspection of the OHC soma shows a continued uptake of the sensor indicating continued MET current (Fig. S6D; bottom panel). Hereto plotting the bundle lifetime against soma intensity provides a strong correlation (r^2^ = 0.84 in the example provided, Fig. S6F). We propose the remaining MET current is due to TMC2 not downregulating at the same rate in *Tmc1^-/-^* as in the *Tmc1^-/+^*.

We also investigated AnV labeling in the *Tmc1^-/-^* mice at P10 (Fig. S7). At P10, the littermate controls showed a range of AnV labeling that was significantly greater than in the *Tmc1^-/-^* which was also significantly greater than in the *Tmc1^-/-^ Tmc2^-/-^* double KO (raw images shown in Fig. S7A and summarized in B). Fig. S7B (bottom panel) overlays AnV and lifetime measurements showing that higher lifetimes and lower AnV labeling correlated with a reduction in the presence of TMC molecules. Both IHC and OHC showed strong correlations between parameters but were different from each other (Fig. S7C).

To further probe MET components, we used the homozygous spinner *Tmie^-/-^* mice^39^ which lack TMIE, a subunit contributing to the channel pore and required for localization of TMC1 and TMC2 to the hair bundle^32,40,41^ and the *Tmc2^-/-^* mice where the onset of MET is delayed until TMC1 expression begins^50^ (Fig. 5). At P10, *Tmie^-/-^* mice had higher lifetimes (*t*-test, *p<0.001*) for the hair bundles and minimal uptake of the sensor in the soma compared to littermate controls, *Tmie^+/-^*, and WT mice (Fig. 5A-C), confirming the need for MET for both the dye uptake and the bundle membrane properties to become less viscous. No significant difference was observed between the PC soma of the *Tmie^-/-^* and the *Tmie^-/+^* and WT mice suggesting that the effect on the membrane viscosity is again specific to the HCs. Hair bundles for both IHCs and OHCs showed uniformly high lifetimes that represent more than an order of magnitude change in effective viscosity (see dye calibration Sup Fig.1). For *Tmc2^-/-^* mice, hair bundle membrane lifetime, measured at the base between P1-P4, was high (*t*-test, *p<0.001*) at P1 compared to littermate controls for both IHCs and OHCs (Fig. 5D-F). By P4, lifetime measurements were comparable between the *Tmc2^-/-^* and *Tmc2^-/+^*, consistent with TMC1 expression compensating for the loss of TMC2. These data are consistent with both TMC1 and TMC2 reducing stereocilia lifetime as predicted by these having scramblase activity. The TMC2^-/-^ data is in contrast with previous reports in hair cells^44^.

**Figure 5:**
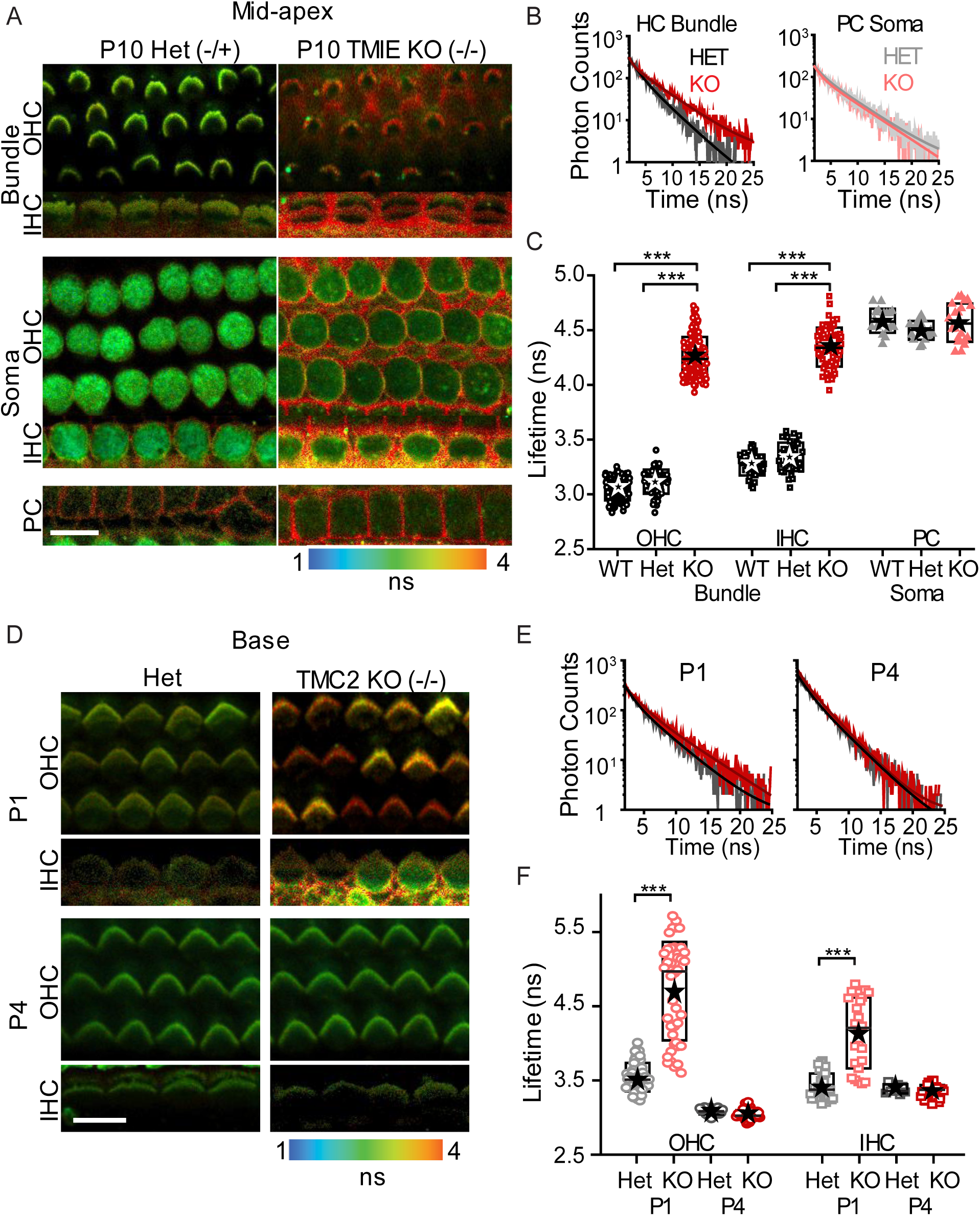
Membrane viscosity is significantly high in mice lacking TMIE and TMC2. A) FLIM images of P10 mid-apical *Tmie*^−/−^ and *Tmie*^−/^ hair bundles (top rows) and soma (bottom row). B) Example fluorescence decay curves comparing the HC bundles and PC soma of mutant mice and the littermate controls. C) Quantification of the mean lifetime from mutants, littermate controls and wild-type mice. D) FLIM images of P1 and P4 basal *Tmc2^-/-^* and *Tmc2^-/+^* mice hair bundles with example fluorescence decay curves in E). F) The mean lifetime from the *Tmc2^-/-^*and *Tmc2^-/+^* mice hair bundles at P1 and P4. Boxes in C and F represent the SD, and the star symbol indicates the mean and the line indicates the median. Each data point corresponds to a hair bundle or a cell (for soma). \*\*\**p* < 0.001. Scale bar = 10 μm.

### MET channel blocking increases stereocilia membrane viscosity

Are the changes in membrane effective viscosity dynamic or established during development? It is possible that membrane properties co-vary with expression of TMCs during development and that the genetic manipulations interfere with the natural maturation process. To test this possibility, we used two known MET channel blockers, tubocurarine (1 mM) and amiloride (1 mM). Curare is an open channel blocker, while amiloride is a permeable channel blocker thought to bind somewhere within the channel pore^54–58^. Fig. 6A shows FLIM images of a 30-minute time course where amiloride was continuously applied. Lifetime initially went up, reached a peak near 7 minutes and then reduced to near baseline by 30 mins. A summary across preparations is presented in Fig. 6B showing the reproducibility of the response. Fig. 6C provides early time point examples of the raw lifetime decay curves for control, curare and amiloride and Fig. 6D shows early and late FLIM images for each compound with Fig. 6E providing a summary across preparations and cells. Both compounds evoked an initial increase in lifetime followed by a slower recovery to baseline levels in the face of continued drug application. The initial increase in lifetime suggests that open MET channels are important for maintaining lower stereocilia membrane lifetimes. It also suggests that membrane lifetime is being dynamically regulated so that when the channels are blocked, effective viscosity increases and when channels are unblocked (and open) effective viscosity reduces. The secondary reduction in viscosity suggests an additional feedback mechanism is involved.

**Figure 6:**
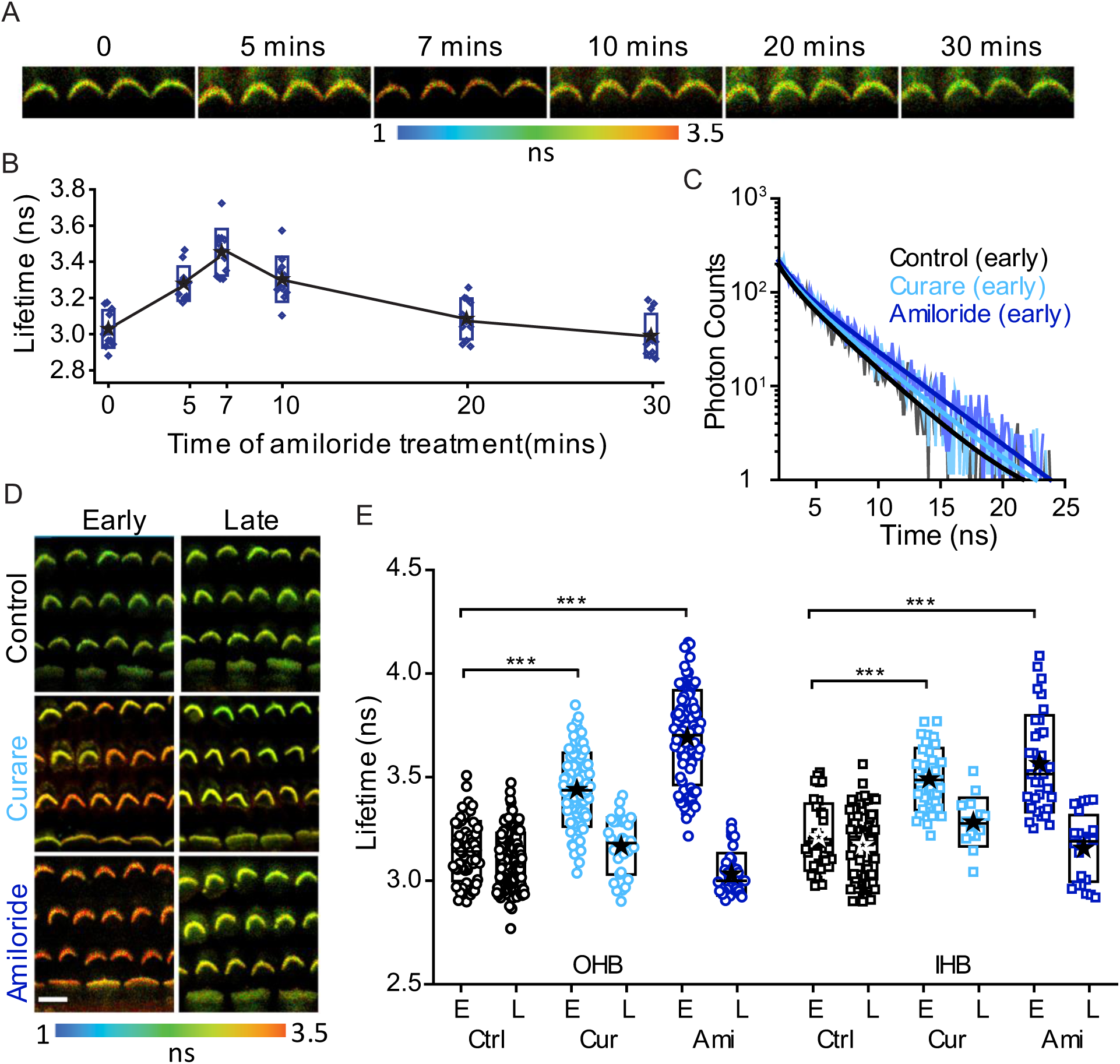
MET channel blocking increases stereocilia membrane viscosity. A) FLIM images of a 30-minute time course where amiloride was continuously applied. B) Quantification of the mean lifetime across samples from hair bundles during amiloride treatment. C) Example lifetime decay curves for control, curare (1 mM) and amiloride (1 mM) from early time point (∼10 mins). D) Early (∼10 mins) and late (∼30 mins) FLIM images of hair bundles for each compound. E) Summary of mean lifetime from hair bundles across preparations and cells for early and late time points from control, curare and amiloride treatments. Boxes in E represent the SD, and the star symbol indicates the mean and the line indicates the median. \*\*\**p* < 0.001. Scale bar = 10 μm.

We also investigated the AnV labelling with curare treatment (Fig. S8). Curare treatment resulted in an increase in the AnV labelling at both early and late time points in both OHC and IHC bundles (*t*-test, *p<0.001*). Thus, PS externalization could be uncoupled from the state of the channel again implying a second scramblase mechanism.

The increase in lifetime with channel block implies dynamic regulation of membrane properties likely involving the presence of a flippase. Given that our earliest measurements show high effective viscosity, there would need to be an early mechanism establishing this high effective viscosity. We blocked MET current at P5, the earliest time point in the mid-apex where membrane viscosity had reduced to determine if the underlying regulation of viscosity was similar. (Fig. S9). Hereto we found an increase in effective viscosity (*t*-test, *p<0.001*) suggesting the membrane was being dynamically regulated, likely increasing membrane order via flippase activity.

### Increase in stereocilia membrane viscosity is independent of MET current and Ca^2+^ entry

To this point both the increases and decreases in lifetime measured could be indirect results of altering mechanotransduction. Current through the MET channel regulates the membrane potential, such that blocking the channel hyperpolarizes the cell potentially triggering a voltage dependent effect (turning off scramblase activity or activating flippases or floppases). Ca^2+^ through the MET channel could be driving some secondary effect (like above) or the blocker when in the pore blocks both MET current and TMC scramblase activity which unmasks an independent flippase activity that increases lifetime. To directly address these potential processes, we took advantage of known MET channel pore properties for example that the channel shows anomalous mole fraction behavior where ions interact within the pore to alter current flow^55,69,70^. Fig. 7A summarizes the manipulations used schematically and presents corresponding stereocilia lifetime images. In our control extracellular solution of Na^+^ as the monovalent ion and Ca^2+^ at 2 mM concentrations, there are large MET currents, despite Ca^2+^ blocking more than 50% of the current and carrying about 60% of the current. When Na^+^ is replaced with an impermeant monovalent ion TRIS^+^, the total current is reduced by more than 50% and Ca^2+^ current reduces because without a permeant monovalent ion, Ca^2+^ inhibits its own permeation^70^. Thus, there is a large total reduction in MET current and a modest reduction in Ca^2+^ entry. Hair cells are expected to hyperpolarize. This manipulation resulted in no statistical difference in membrane lifetime suggesting that neither current nor voltage were responsible for the effect on membrane seen with the channel blocker. Summary data are shown in Fig. 7B. Lowering external Ca^2+^ increases the total MET current by reducing the Ca^2+^ block of the channel and increases the resting open probability of the channel by a slow adaptation process^70–72^. These changes are expected to depolarize hair cells. Fig. S10 shows that lowering Ca^2+^ to 20 μM while maintaining Na^+^ as the monovalent resulted in a decrease in lifetime. This decrease in lifetime could be due to any of the factors described above. To isolate the MET channel being open from both Ca^2+^ and depolarization, we used a TRIS^+^ and 20 μM Ca^2+^ solution. In this case, MET channels will open, however there will be significantly less total current and even less permeation by Ca^2+^, resulting in the hair cells hyperpolarizing despite the channels being open^70^. In this case, stereocilia lifetime reduced (*t*-test, *p<0.001*) for both IHCs and OHCs (Fig. 7). Thus, neither Ca^2+^ nor current (or membrane potential) are responsible for the faster lifetime, rather the increased open probability of the channel likely reflects an increased scramblase activity. Adding amiloride to the TRIS^+^ and 20 μM Ca^2+^ solution confirms our result as lifetime increased in the presence of the blocker despite their being no change in current or calcium entry (*t*-test, *p<0.001*) (Fig. 7). Given that under these circumstances there is little effective current or Ca^2+^ permeation, the increased viscosity is most likely ascribed to the channel blockers directly inhibiting the scramblase activity. The relative change in lifetime between control solution and amiloride application and the TRIS^+^ + 20 μM Ca^2+^ solution and TRIS^+^ + 20 μM Ca^2+^ + amiloride are comparable, suggesting that the underlying mechanism is the same and independent of current and Ca^2+^.

**Figure 7:**
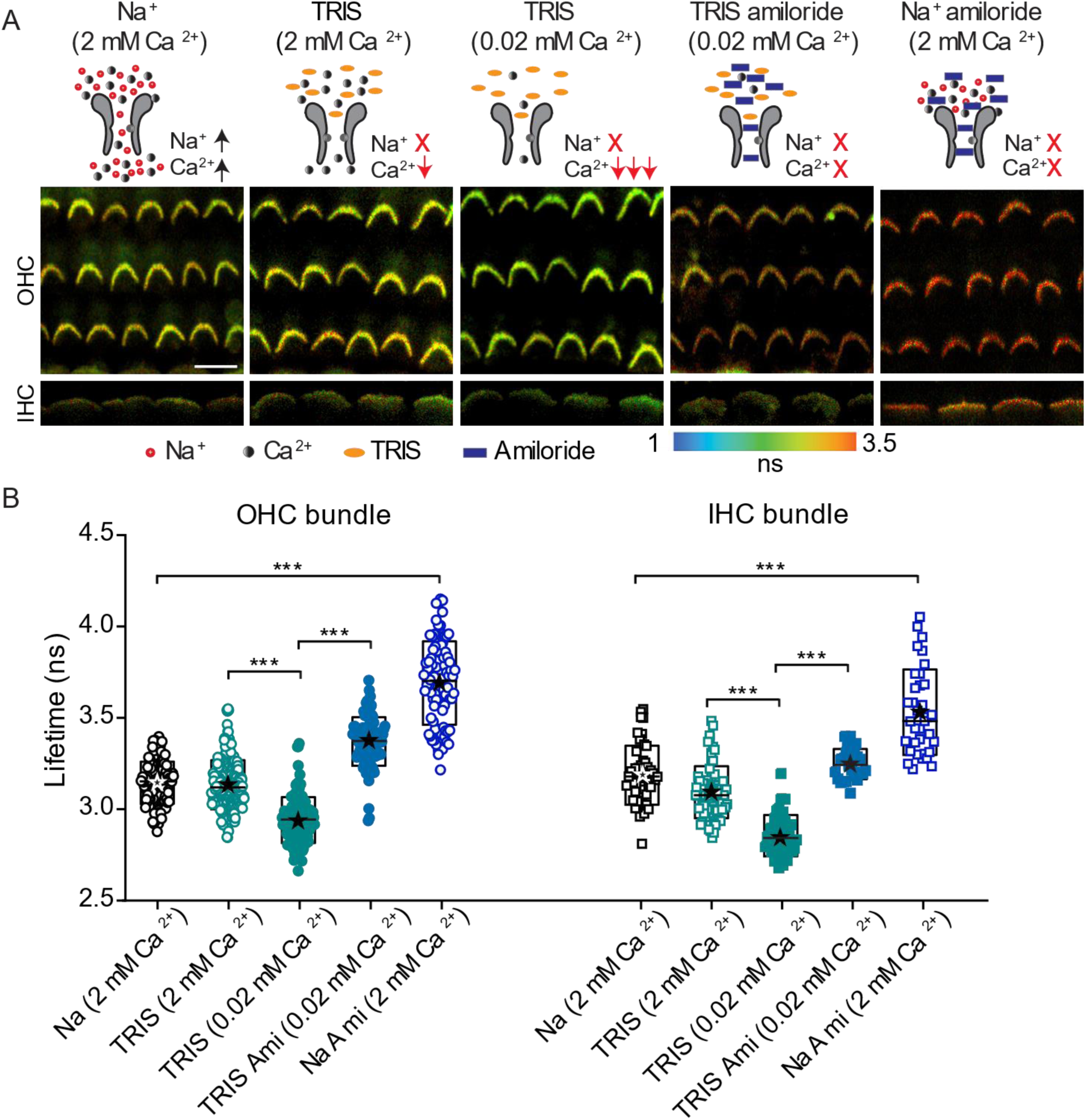
Increase in stereocilia membrane viscosity is independent of MET current and Ca^2+^ entry. A) FLIM images from P10 rat mid-apical organ of Corti with different monovalent ions (Na^+^ vs TRIS^+^), Ca^2+^ concentrations (2 mM vs 0.02 mM) with presence or absence of 1 mM amiloride. The manipulations are represented schematically above each representative FLIM image. B) Summary plot showing the mean lifetime measured from the hair bundles from each manipulation. Each manipulation was performed in at least 4 animals. Boxes in B represent the SD, and the star symbol indicates the mean and the line indicates the median. \*\*\**p* < 0.001. Scale bar = 10 μm.

## Discussion

Our data provides the first evidence that stereocilia membrane effective viscosity is regulated by scramblase activity associated with the MET complex and a second process responsible for increasing membrane effective viscosity that counters the scramblase activity. Several pieces of evidence target the MET complex driving the reduction in membrane viscosity (i.e. lowering lifetime, Fig. 8A). There was a strong correlation between developmental onset of MET current, and stereocilia membrane lifetime. MET channel mutants, including TMC1, TMC2, TMC1/2 double KO and TMIE, showed higher lifetimes as compared to their littermate controls. MET channel blockers increased membrane lifetime under conditions where the channels were open, but current was not flowing; thus, ruling out the effects of current, voltage or Ca^2+^ and providing direct evidence that the MET channels provide scramblase activity that could be inhibited by these same channel blockers. The concomitant increase in effective viscosity in the presence of the channel blocker provides evidence for an independent mechanism being responsible for increasing membrane effective viscosity.

**Figure 8:**
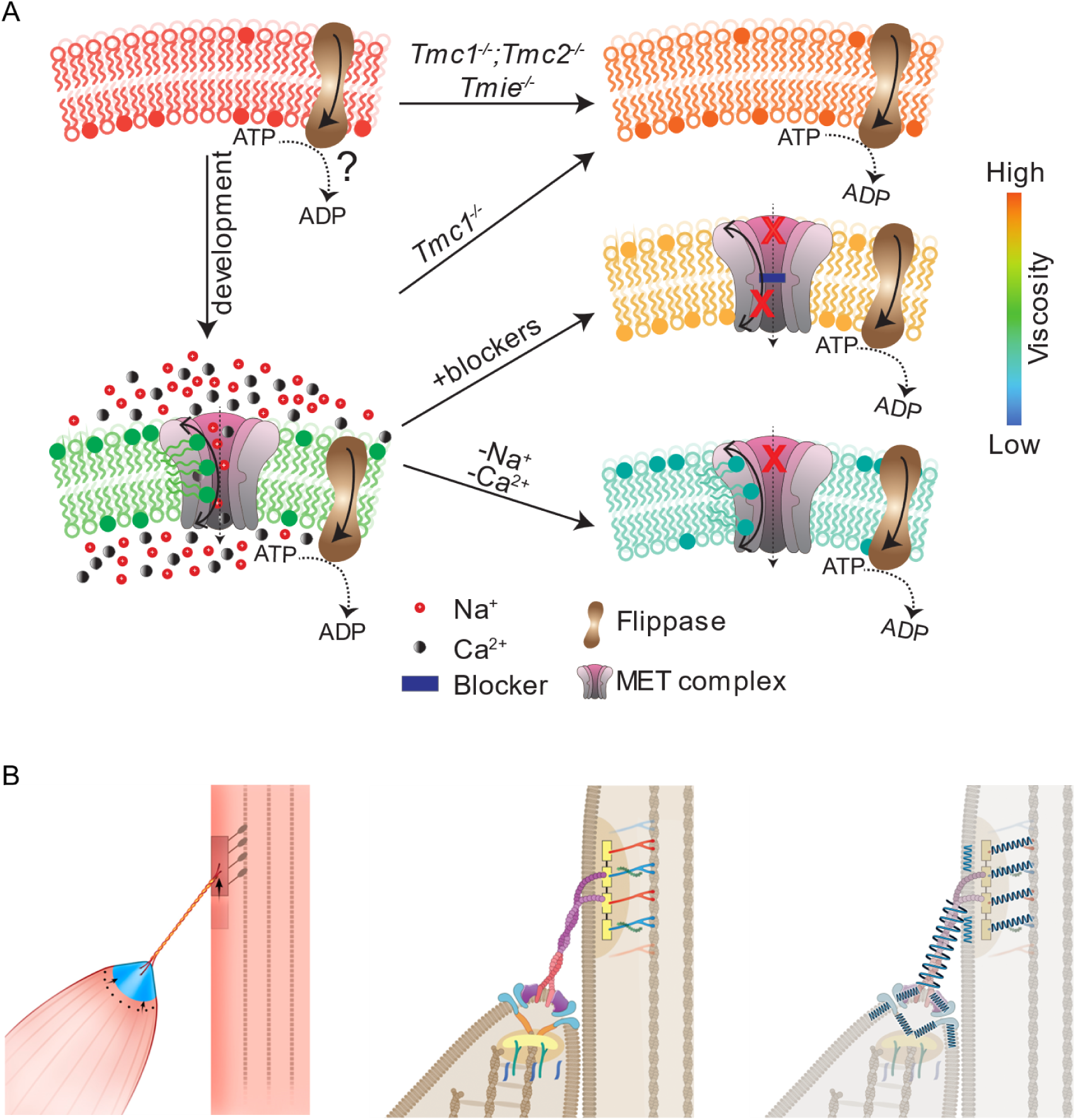
Dynamic modulation of auditory hair cell stereocilia membrane mechanics by the MET complex. A) Developmental, genetic, electrophysiological and pharmacological data demonstrate that both TMC1 and TMC2 are mechanosensitive scramblases as activity depends on MET channel being open. Blockage of scramblase activity unmasks a second process, potentially based on flippases, that increases effective membrane viscosity independent of MET machinery. B) Depending on how the various components of the MET machinery are mechanically coupled together, the impact of the plasma membrane on the force translation pathway will be determined by whether the membrane is more in series or in parallel.

### Active lipid transport

This study suggests there is a dynamic interplay between scramblase activity sensitive to the MET open probability and an independent mechanism serving to increase membrane effective viscosity that rapidly modulates membrane mechanical properties. We hypothesize flippase/floppase activity is involved from the very high effective membrane viscosity prior to the onset of mechanotransduction, in the mutant mice that eliminate the presence of the MET complex or the TMC components of the complex and with blockers of the MET channel resulting in increased effective viscosity. The high viscosity could also be attributed to lipid diffusion when the scramblase is blocked, different lipid compositions being present during development, membrane interaction with underlying cytoskeletal structures, mechanotransduction machinery and/or active lipid transport (like flippases)^73,74^. Significant developmental changes in membrane properties have been previously reported in other systems and associated with lipid composition^75–77^. However, the highest lifetime reported in our study is like that reported for model membranes made entirely of sphingomyelin and cholesterol, one of the most viscous lipid combinations known and a composition unlikely to be found in stereocilia at any age. Additionally, the lipid composition cannot account for the increased effective viscosity following MET channel blocker application as early as postnatal day 5. The increased effective viscosity is consistent with an enhanced membrane order typically achieved by active transport via flippase or floppase activity. If effective viscosity were not dynamic, the channel blocking experiments would predict no viscosity change. The stereociliary membrane specifically expresses ATP8B1 flippase that transports PS from the outer to the inner membrane. Morphological and functional degeneration of the hair bundles due to the mutations of ATP8B1 and ATP8A2 indicate the importance of this transport mechanism involved in maintaining the stereocilia^78,79^. However, recent ongoing studies indicate that ATP8B1 flippase starts expressing in the stereocilia from post-natal day 7. This timing is consistent with our Annexin labeling but suggests an additional pathway, perhaps ATP8A2 which is present in hair bundles as early as P0. Previous work using AnV labeling suggest that TMC1 and not TMC2 is required for scramblase activity in mammalian hair cells^44^. Our work suggests that both TMC1 and TMC2 are involved in stereocilia membrane regulation. Both developmental data showing an early decrease in effective viscosity when TMC2 is being expressed and the TMC1^-/-^ mouse showing a decrease in effective viscosity are consistent with TMC2 having scramblase activity. Previous work also suggested that Ca^2+^ buffering was activating scramblase activity. Our study shows that blocking the channel leads to an initial increased lifetime, followed by a recovery and decrease in lifetime. The latter is consistent with the previous reports and our AnV labelling also agrees with previous reports. The viscosity sensor demonstrates though that PS labeling is limited in that it is not clear that all scramblases scramble PS equally and scramblases scramble more than PS. Our data shows a restoration of scramblase activity in the face of channel blockers implying non-MET channel scramblases being activated.

### Scramblase Speed

Are the molecules associated with the MET channel capable of altering the effective membrane viscosity of the entire stereocilia, including row 1 stereocilia where TMC molecules have not been identified? Recent evidence of TMEM16F scramblase rates suggests up to 45,000 lipids per second^80^. A stereocilium modelled as a cylinder of 0.4 µm diameter and a height of 4 µm^81^ produces a surface area of ∼5.3 µm^2^. Previous reports indicate an average surface area for each lipid molecule of ∼0.5 nm^2^ ^82–84^, and assuming two functional channels in a stereocilium^5,85^, we can estimate it would take 3.9 mins for the channel to turnover its entire stereocilium membrane. It has been suggested that there are more TMC molecules per stereocilia than account for 2 MET channels and if these contribute to scrambling, the time to turn over the stereocilia would be much shorter^62^. Finally, this rough calculation does not include membrane proteins that would reduce the lipid concentration and further speed up membrane turnover.

### Functional Relevance

The present work identifies a dynamic regulation of stereocilia membrane mechanical properties that is regulated by mechanically-sensitive scramblase activity associated with the MET complex and a mechanism increasing membrane effective viscosity (potentially an ATP-dependent flippase). Given that hair cell MET operates at the highest known rates of any mechanoreceptor system and has both measured and calculated sensitivities at molecular dimensions, a simple idea is that reducing membrane viscosity will lower the energy for channel gating which may enhance both the frequency and sensitivity response^1,86,87^.

Our data suggests that the membrane is being actively regulated and at least the scramblase activity is sensitive to the MET channel open state and so is also mechanically sensitive. When the MET channel opens effective membrane viscosity will reduce. If reducing membrane viscosity leads to channel closure, one can envision how this might contribute to the fast adaptation response, where local viscosity changes could happen in the milli to sub millisecond time frame. If lowering viscosity opens channels, then one might envision cooperativity between channels like what has been predicted computationally^24^. Direct measurements are needed to sort this out.

Another way to envision the membrane impact on the MET process is to consider the complexity of the molecular coupling of the machinery (Fig. 8). Previous studies have reported the existence of multiple molecular interactions between the MET channel components and the lower part of the tip link, PCDH15. For example, TMC1, TMC2, TMIE, LHFPL5, LOXHD1 and Myo15 can interact with PCDH15 and each of these molecules also interacts directly with the membrane^9,88–90^. Depending on how this machinery is wired together, the plasma membrane might be considered in series and/or in parallel with the force translation pathway. Fig. 8 depicts that the membrane tenting when the tip link is pulled will interact with the various components of the MET machinery differently depending on the mechanical coupling and that this could be direct or indirect. The relative contribution of the plasma membrane will depend in part on the mechanical coupling between these molecules, but the impact of membrane viscosity changes will be determined by whether the membrane is more in series or parallel. If in parallel, a reduction in effective membrane viscosity would be expected to shift more of the force stimulation onto the channel, allowing it to open faster and in response to a smaller total force stimulus. Increasing the effective viscosity would result in the channels receiving less of the total force, requiring larger stimulations for channel gating and resulting in slower responses. If in series, force would be the same, however motion would change with the lower effective viscosity resulting in larger motion (i.e. spring extension).

And finally, as the scramblase is using energy likely provided by the flippase activity that establishes an electro-lipid gradient across the bilayer, it is plausible to imagine this system contributing to hair bundle amplification^1^. Thus, this newly developed membrane regulation that is established by the interplay of flippases/floppases and scramblases that are also mechanically sensitive ion channels provides a new framework from which to evaluate old and new data about hair bundle mechanics and MET channel properties.

Our data supports the hypothesis that the MET channel complex has scramblase activity and posits that this activity is fundamental to mechanosensitivity. Given that there are likely direct impacts of membrane mechanics on ion-permeation and vice versa, a reevaluation of many of the known TMC mutations on membrane mechanical properties is warranted, as a means of delineating between residues impacting scramblase vs ion permeation. Also, given that PS is a major negatively charged lipid that can be scrambled at high rates, it is possible that permeation estimates are impacted by changes in scramblase activity. It is also possible that some larger known molecules that permeate MET channels, likely FM1-43, aminoglycosides are taking advantage of scrambling pathways rather than canonical ion channel permeation^59,91–94^.

In conclusion, our data provides the first evidence of membrane scramblase activity impacting stereocilia effective viscosity and that the scramblase activity is generated by TMC components of the MET channel complex. The interplay between mechanosensitivity, scramblase and flippase/floppase activity provides a new framework from which to evaluate long-standing MET phenomena like channel gating and permeation, adaptation, gating compliance and even hair bundle amplification. It is likely that the dynamic membrane regulation is not unique to hair cell stereocilia but may in fact be a yet unrecognized partner in regulation of excitable cells.

## STAR Methods

### Sample Preparation

#### Cochlear Explants

All animal experiments were approved by the Stanford University Administration Panel on Laboratory Animal Care (APLAC #14345). Rodents (rats and mice) of both sexes were used in the experiments. The wild-type (WT) C57BL6/J mice and Sprague-Dawley rats were purchased from Charles River Laboratories. *Tmie^KO^* ^39^ (B6.B(CBA)-*Tmie^sr^*/J, JAX 000543) was obtained from JAX, *Tmc1^KO^* ^50^ (B6.129-*Tmc1^tm^*^1^*^.1Ajg^*/J, JAX 019146, backcrossed on C57Bl6/J) from Dr. Beurg and Dr. Fettiplace (Wisconsin University), *Tmc2^KO^* from Dr. Angela Ballesteros^44^ and *Tmc1^HA^* were obtained from Dr. Cunningham and Dr. Muller (J. Hopkins University)^32^. For mice, heterozygous littermates served as controls. Experiments were performed blinded to genotype. The mice were genotyped as previously described.

Animals were sacrificed by decapitation and cochlear turn of isolated organ of Corti was dissected from pups at postnatal day 1 (P1) to P10 and placed in a recording chamber as described previously^5^. Briefly, temporal bones were removed from the skull and cochlear tissue was placed in extracellular solution containing (in mM): 142 NaCl, 2 Cl, 2 CaCl^2^, 1 MgCl^2^, 10 HEPES, 6 Glucose, 2 Ascorbate /Pyruvate, 2 Creatine monohydrate, at pH 7.4 and a final osmolality of 304 - 307 mOsm. After removing the semicircular canal and vestibular organs, the cochlear bone was removed to expose the organ of Corti. The modiolus, Reissner’s membrane, and the tectorial membrane were sequentially removed. The tissue was then incubated with 10 µM of BODIPY (boron dipyrrin) 1c in extracellular solution at room temperature (RT) for 6 mins, protected from light. BODIPY 1c stock solution (100 mM) was prepared in 100% ethanol and stored at -20^°^C. Working dye solution was made daily by diluting in extracellular solution to a final BODIPY 1c concentration of 10 µM in 0.01% ethanol and kept at room temperature (RT; 19 - 21°C), protected from light.

The tissue was then transferred to the recording chamber with dye-free extracellular solution and held in place with single strands of dental floss while ensuring that inner hair cell (IHC) bundles were oriented vertically. During the experiment, tissue was perfused with extracellular solution at rate of 0.3 ml/min maintained at RT.

### Fluorescence Lifetime Imaging (FLIM)

FLIM experiments were carried out using an upright Leica TCS-SP8 FALCON (FAst Lifetime CONtrast) confocal microscope using LAS X version 3.5.7 and LAS X FLIM/FCS 3.5.6 softwares (Leica Microsystems GmbH, Germany) and an HC APO 20x / 1.0 NA water dipping objective (Leica Microsystems GmbH, Germany). The excitation source was a pulsed white light laser (WLL) with an acoustic-optical beam splitter (AOBS). BODIPY 1c was excited at 480 nm and the pulse repetition frequency of the laser was set to 40 MHz. Two sensitive hybrid detectors (SMD, Leica Microsystems CMS GmbH, Germany) allowed for photon detection from 490-560 nm and 600-670 nm. A frame accumulation of 50 was set to allow better photon statistics. The laser intensity was set to keep the photon count of >= 1 per laser pulse to avoid photon “pile up” that leads to shortening of the fluorescence lifetime (Suhling et al., 2012). Z-stacks were captured at 1 um step interval.

### FLIM Analysis

All FLIM images were analyzed using LAS X FLIM/FCS version 3.5.6 software. Each ROI consisted of a hair bundle on a single z-plane. Pixels were binned to have at least 300 photon counts at the peak of the decay for reliable analysis. The fluorescence decay curves were fitted using n-exponential reconvolution with IRF (instrument response function) model. The function of this model is

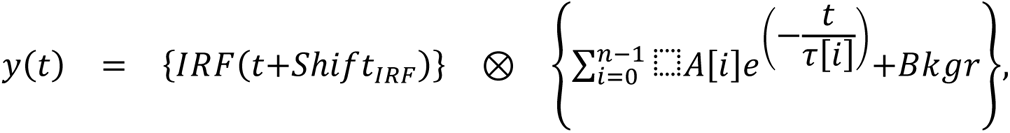

where A is the amplitudes, τ is the lifetimes, n is the number of exponential components, 𝐵𝑘𝑔𝑟 is the tail offset (correction for background), and 𝑆ℎ𝑖𝑓𝑡_𝐼𝑅𝐹_ is the IRF shift (correction for IRF displacement). For the hair bundles, the decay curves were best fitted by biexponential model with the goodness of fit parameter (χ^2^) >1. The average of the two intensity-weighted lifetime components were calculated by

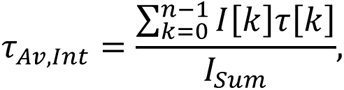

where Isum is the sum of the fluorescence intensity of all components. The intensity-weighted average lifetime represents the mean photon arrival time and hence we report this value in this paper for all the experiments.

### Drug Treatment

*MET channel blockers:* In a subset of experiments, during and post BODIPY 1c staining, the tissue was incubated in 1 mM tubocurarine (93750, Sigma) or 1mM amiloride (0890, TOCRIS) to block MET channel.

### Electrophysiology

In several experiments, FLIM was combined with whole-cell patch clamping to measure MET and membrane viscosity from the same hair bundle. The tissue was viewed in brightfield using a custom built fixed-stage upright BX 51WI (Olympus) microscope with a digital Rolera-XR camera (QImaging). Orientation of pipettes and access to the hair cells were as previously described^95^. Whole-cell patch clamp recordings were obtained using thick-walled borosilicate patch pipettes (WPI) pulled on a P97 micropipette puller (Sutter Instruments) to 1-2 μm inner diameters (3-3.5 MΩ). The internal solution contained (in mM): 116 CsCl, 3.5 MgCl^2^, 3.5 ATP, 5 creatine phosphate, 1 Tetracesium BAPTA, 10 HEPES, and 20 ascorbic acid, pH of 7.2 and osmolality of 285 - 290 mOsm. Extracellular solution was delivered through a bath solution and also through an apical pipette of tip size ∼200 μm^20^. The apical perfusion pipette prevented internal solution from reaching BAPTA sensitive hair bundles.

Outer hair cells were voltage-clamped cells at -80 m and adjusted offline for liquid junction potential (4 m). Mechanically elicited currents were low pass filtered at 10 kHz, with an Axopatch 200b patch clamp amplifier (Axon Instruments), sampled at 100 kHz using Personal DAQ3000 (IOtech), and recorded with jClamp (SciSoft). Uncompensated series resistance (Rs) was between 6 and 10 MΩ.

#### Mechanical Stimulation

Hair bundles were deflected with a custom-built fluid jet system as previously described^64^. Thin-walled borosilicate pipettes (10 µm tip diameter) were filled with extracellular solution and positioned within 5 μm of the voltage-clamped hair bundle. The fluid jet was driven with a 50 Hz sinusoidal wave using a piezo electric disc bender whose input was filtered with an 8-pole Bessel filter (Frequency Devices) at 1 kHz before and then amplified by a high voltage/high current custom-built amplifier.

### Immunofluorescence staining and imaging

In a few experiments, following the FLIM experiment, some cochlear turns were stained for TMC1 HA. The dissected sample was fixed in 4% PFA in extracellular solution for 40 mins at RT. The fixed samples were washed with PBS 3 times and transferred to a glass well plate with PBS containing 0.05% Triton X-100, permeabilized for 40 mins at RT. After permeabilization, the samples were blocked in PBS with 0.05% Tween 20 (PBST) containing 4% bovine-serum albumin Fraction (BSA) overnight at 4℃. The tissues were then incubated with primary antibody (Rabbit anti-HA, CST #3724, 1:500) in PBST with 1% BSA (incubation buffer) overnight at 4 ℃. The samples were then washed 4 times, 5-10 minutes per wash, in incubation buffer at RT. Then the tissues were incubated with fluorescent dye conjugated secondary antibodies (Donkey anti-Rabbit IgG with Alexa Fluor 488, Invitrogen A21206, 1:500) in incubation buffer at RT for 1-2 hours. After 1 wash (5-10 min) with incubation buffer, the samples were incubated with fluorescent dye conjugated Phalloidin (Alexa Fluor 568 Phalloidin, Invitrogen A12380, 1:500) in incubation buffer at RT for 25 minutes. Then the samples were washed 3 times, 5-10 minutes per wash, with incubation buffer. The glass well plate was on a horizontal shaker with a 60-rpm speed during permeabilization, incubation and washing steps. Each experiment contained at least one parallel stained cochlea sample from a similar age mouse without any tag, as background control.

After washing, the tissue was transferred to the recording chamber with PBS and held in place with single strands of dental floss while ensuring that the hair bundles were oriented vertically. Z-stacks were captured using the Lightning super-resolution mode of an upright Leica TCS-SP8 confocal microscope with an HC PL APO 63x / 1.20 water immersion objective lens and LAS X 3.5.7 software. TMC1 HA and phalloidin were excited using a white light laser that can be tuned for wavelengths between 470 nm – 670 nm. The image acquisition parameters were determined as the best X*Y and Z axis resolution possible for the shortest wavelength used.

### Data Analysis

We used ImageJ for the intensity measurements of BODIPY 1c in the hair cell soma, and the hair bundle intensities measurements of An and TMC1 HA. Whole cell currents were visualized and analyzed using jClamp (SciSoft) and OriginPro 2018 (OriginLabs). Graphs were generated with OriginPro 2018 (OriginLabs) and Adobe Illustrator CS6 (Adobe). All statistical analyses used two-sample Student’s t test performed using OriginPro 2018 (OriginLabs). Significance (*p* values) are * *p* < 0.05, ** *p* < 0.01, *** *p* < 0.001. Data are presented as mean ± SD.

## Acknowledgements

This work was supported by National Institute on Deafness and other Communication Disorders (NIDCD) grant RO1 DC003896 and RO1 DC014658 to A.J.R and R21 ECR DC021027 to S.S.G.

We would like also to thank the Oberndorf family and other SICHL contributors for their support.

## Author Contributions

S.S.G. and A.J.R. designed the experiments, S.S.G. performed the experiments and analyzed the data, S.S.G. and A.J.R. wrote the manuscript.

