## Supplemental for "Dynamic modulation of auditory hair cell stereocilia membrane mechanics by the scrambling mechanotransduction complex"

### Supplementary Information

#### Validation and calibration of BODIPY 1c

BODIPY 1c has been calibrated previously by the lab that designed it (Sherin et al., 2017), but we characterized the 1c that we synthesized for our lab (Nanosyn) under our experimental conditions and with our imaging systems. Artificial lipid vesicles were used as a control to validate dye properties. Liposomes were prepared with either 100% 1,2-dioleoyl-sn-glycero-3-phosphocholine (DOPC, Avanti Polar Lipids) or 70% Egg Sphingomyelin (Egg SM, Avanti Polar Lipids, 860061).and 30% Cholesterol (Ovine Cholesterol, Avanti Polar lipids, 700000). For 100% DOPC vesicles, the solvent of a 0.1 ml aliquot of 10 mg/ml DOPC in chloroform was evaporated in the vacuum to leave a dried lipid film. To prepare SM/Chol vesicles, the solvent containing 70 µl of 10 mM SM and 30 µl of 10 mM of cholesterol in chloroform was dried in the vacuum. Dried lipids were hydrated in the buffer (0.25ml for 100% DOPC and 0.3 ml for SM/Chol) with 100 mM KCl and 10 mM HEPES at pH 7.4 for an hour. The lipid solution was then extruded (Mini Extruder, Avanti Polar Lipids) through a 200 nm polycarbonate membrane 11 times. BODIPY 1c was added to the lipid solution in 1:200 rotor:lipid ratio to prevent dye aggregation and kept at 4°C, away from light. For imaging, the liposomes were mounted on a slide in 0.5% agarose gel (Fisher Scientific) to keep them mechanically stable. Vesicles were prepared on the day of the experiment. As shown previously (Sherin et al., 2017), the highly viscous SM:Chol vesicles showed slower fluorescence decay curves and higher lifetime than pure DOPC vesicles (Fig. S1A-C), thus validating 1c. Importantly, BODIPY 1c showed similar lifetime values in these vesicles as previously reported at 20°C (Sherin et al., 2017).

To perform quantitative measurement of viscosity, the lifetime of BODIPY 1c was calibrated as a function of viscosity using glycerol (Acros Organics, CAS 56-81-5) and methanol (Fisher Chemical, A456-1) mixtures varying from 30:70 to 100:0 (Fig. S1D-F). BODIPY 1c was added to the 2 ml glycerol/methanol mixture at 1:500 rotor:mixture ratio and kept on a rotating vertical shaker (Labquake) at room temperature, away from the light, overnight for efficient mixing. For fluorescence lifetime imaging, the glycerol/methanol mixture was transferred to a glass well plate and imaged using Leica SP8 FALCON system at 20°C.

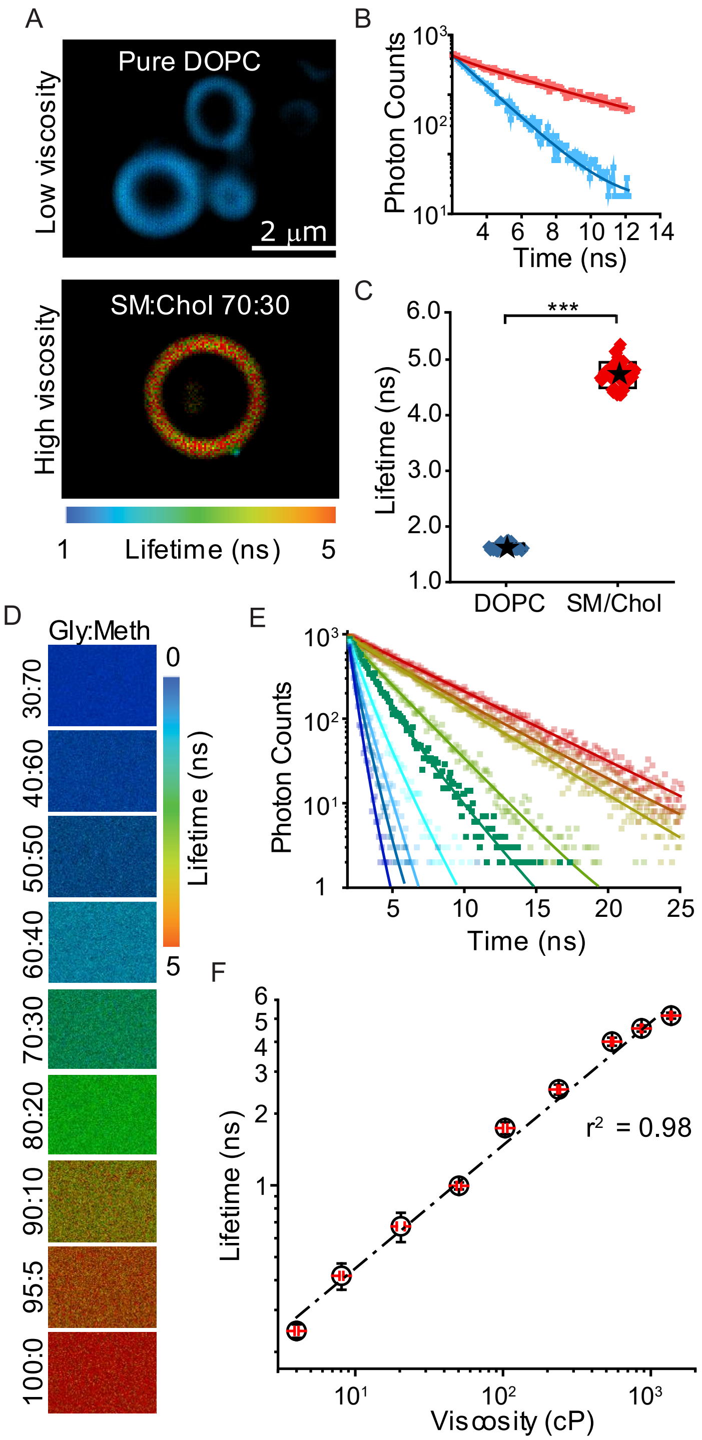


**Supplementary Figure 1:** Validation and calibration of BODIPY 1c in model membranes and glycerol/methanol mixtures. A) FLIM images of BODIPY 1c in pure DOPC (top panel) and SM:Chol 70:30 (bottom panel) vesicles. B) Time-resolved fluorescence decays recorded and C) Lifetimes measured from pure DOPC (blue) and SM:Chol (red) vesicles. D) FLIM images and E) Time-resolved fluorescence decay of BODIPY 1c in glycerol/methanol mixtures of varying composition. F) Fluorescence lifetime vs. viscosity calibration obtained for BODIPY 1c in glycerol/methanol mixtures at 20°C.

Viscosity measurements of the above glycerol/methanol mixtures (20/80 to 100/0) were made with ARES-G2 rheometer (TA Instruments) with APS temperature control system and a 40 mm cone plate geometry. Samples for viscosity measurements were 0.8 ml and measured at shear rate from 10 to 100 1/s for 30s; the flow cycles were run thrice for each mixture. The plate unit was kept at a constant temperature of 20°C with water circulating from a temperature-controlled water bath. A logarithmic plot of the fluorescence lifetime versus the mixture viscosity yielded a straight line (Fig. S1F, $r^2 = 0.98$) that obeys the Förster Hoffman equation (Förster & Hoffmann, 1971). This plot was very similar to that previously generated (Sherin et al., 2017) and served as a calibration graph to convert fluorescence lifetime to viscosity.  We report our data as lifetimes because we cannot calibrate our sensor in the native hair bundle environment where cytoskeletal interactions and membrane proteins might alter the absolute value of viscosity. Our data does suggest that reporting an ‘effective viscosity’ is valid and discussing changes observed with the sensor as impacting viscosity is valid.

#### Determining the BODIPY 1c concentration for cochlea

BODIPY based molecular rotors are characterized by monoexponential fluorescence decays in homogeneous medium and in the absence of aggregates. Hence, the presence of biexponential decays can be either the presence of aggregates or lipid heterogeneities in the membrane. The aggregated species are characterized by a weak emission in the red region from 600-670 nm (Sherin et al., 2017). The aggregates cause quenching of the main emission band from 490-560 nm which renders the lifetime-viscosity calibration curve unstable. We, therefore, compared the decay curves recorded at 490-560 nm and 600-670 nm at a range of BODIPY 1c concentrations in cochlear hair bundles (Fig. S2A) to determine an optimal incubation concentration for cochlear cells that is low enough to avoid dye aggregation and high enough to achieve good staining. If the decay curves from the two wavelength ranges are different, that indicates the presence of dye aggregation. The decay curves recorded from the two spectra for BODIPY 1c in cochlear hair bundles show evidence of dye aggregation at 16 and 12 μM but not at 10 μM (Fig. S2A). Hence, we used a concentration of 8-10 μM for all our subsequent experiments. We also confirmed that the fluorescence lifetime measurements were independent of the dye concentration below 10 μM (Fig. S2B-D).

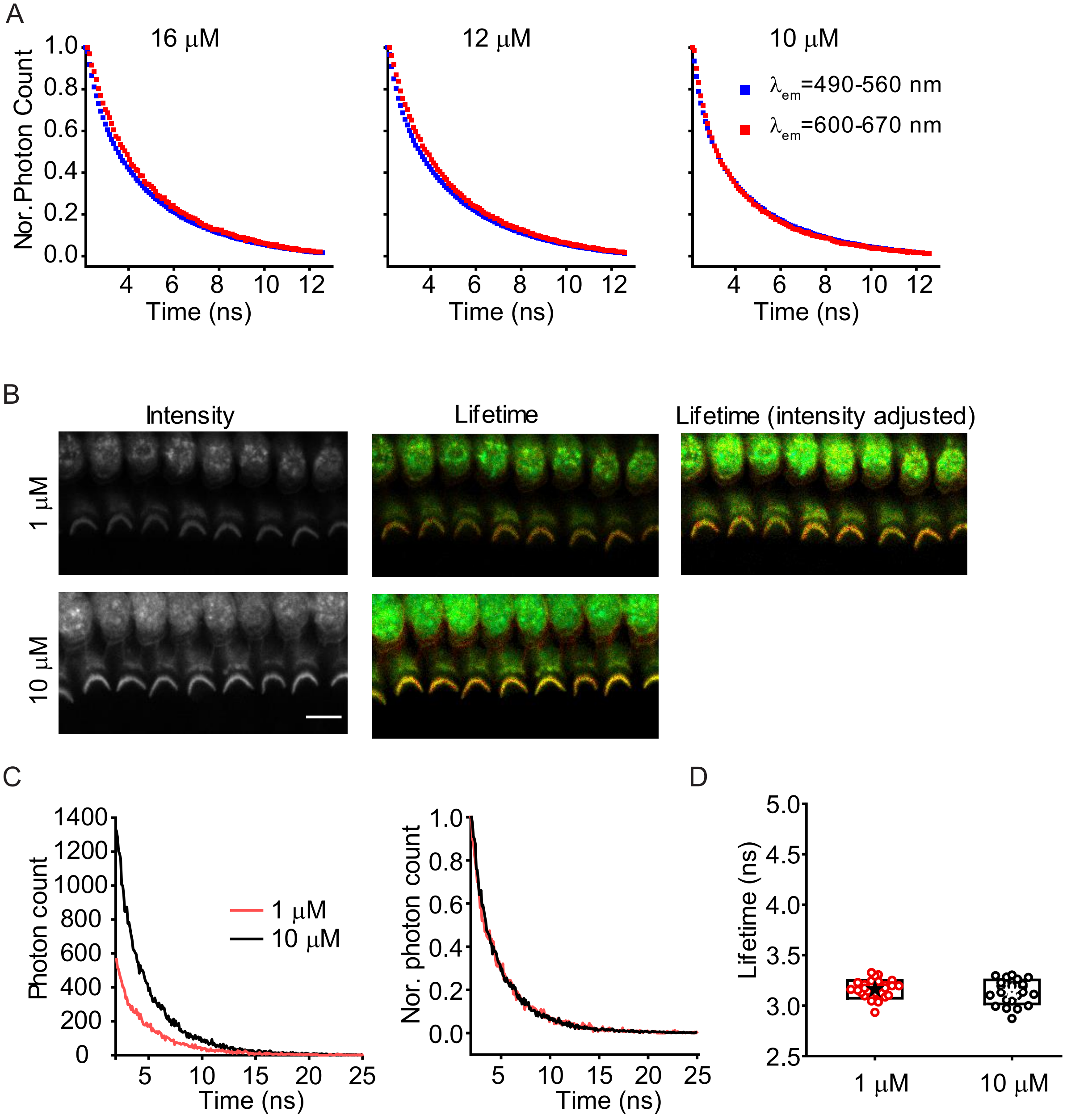

**Supplementary Figure 2:** A) Time-resolved fluorescence decays recorded from P10 rat mid-apical cochlear hair bundles stained with different concentrations of BODIPY 1c (16, 12 and 10 μM) in PBS following excitation at 480 nm and detection in two spectral ranges 490-560 nm (monomers) and 600-670 nm (aggregates). B) Intensity and FLIM images, C) Time-resolved fluorescence decay curves and D) Measure lifetime of P10 rat mid-apical cochlear hair bundles at 1 μM (top row in B, red lines and symbols) and 10 μM (bottom row in B, black lines and symbols) concentrations of BODIPY 1c. Scale bar = 10 μm.

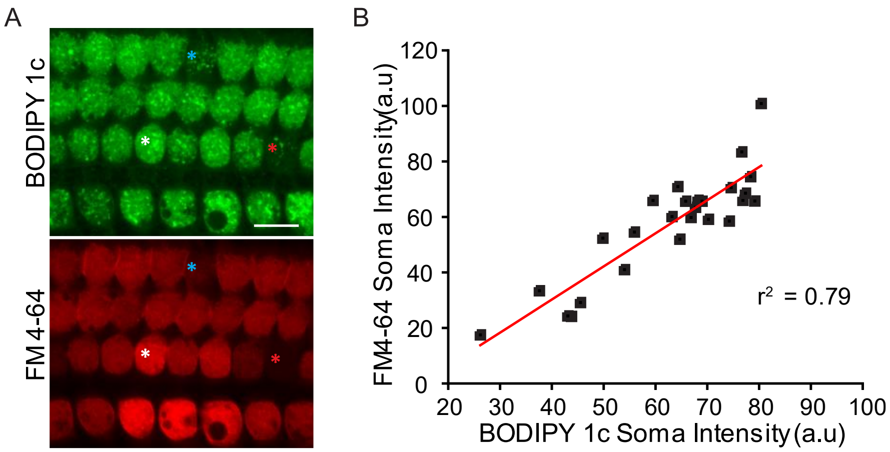


**Supplementary Figure 3:** A) Intensity images of BODIPY 1c (top panel) and FM 4-64 (bottom panel) from the soma of same sample (P10 rat mid-apical turn). Asterisks of different colors are used to highlight a particular cell. B) Plot showing the correlation between the soma intensity of BODIPY 1c and FM 4-64. Scale bar = 10 μm.

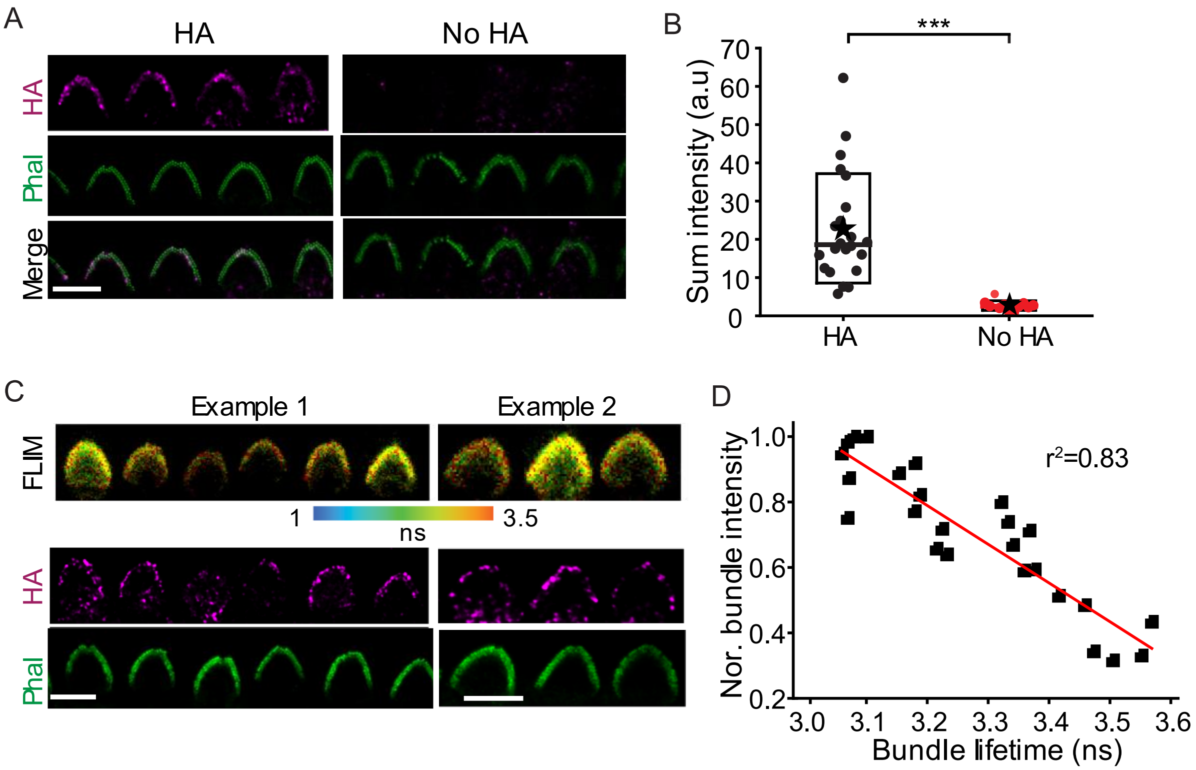


**Supplementary Figure 4:** A) Confocal images of actin-rich stereocilia labelled with phalloidin (green), and with an anti-HA antibody to detect TMC1 in *Tmc1*$^{HA/HA}$ animals. Specific HA staining is detected in *Tmc1*$^{HA/HA}$ mid-apical hair bundles at P4 but not in negative controls without HA. B) Quantification of HA intensity from *Tmc1*$^{HA/HA}$ and negative controls without HA. C) Two example sets showing FLIM and corresponding confocal images of HA and phalloidin. D) Plot showing the correlation ($r^2$= 0.83, $p$ < 0.001) between the normalized sum of HA intensity and the bundle lifetime. Scale bars = 5 μm.

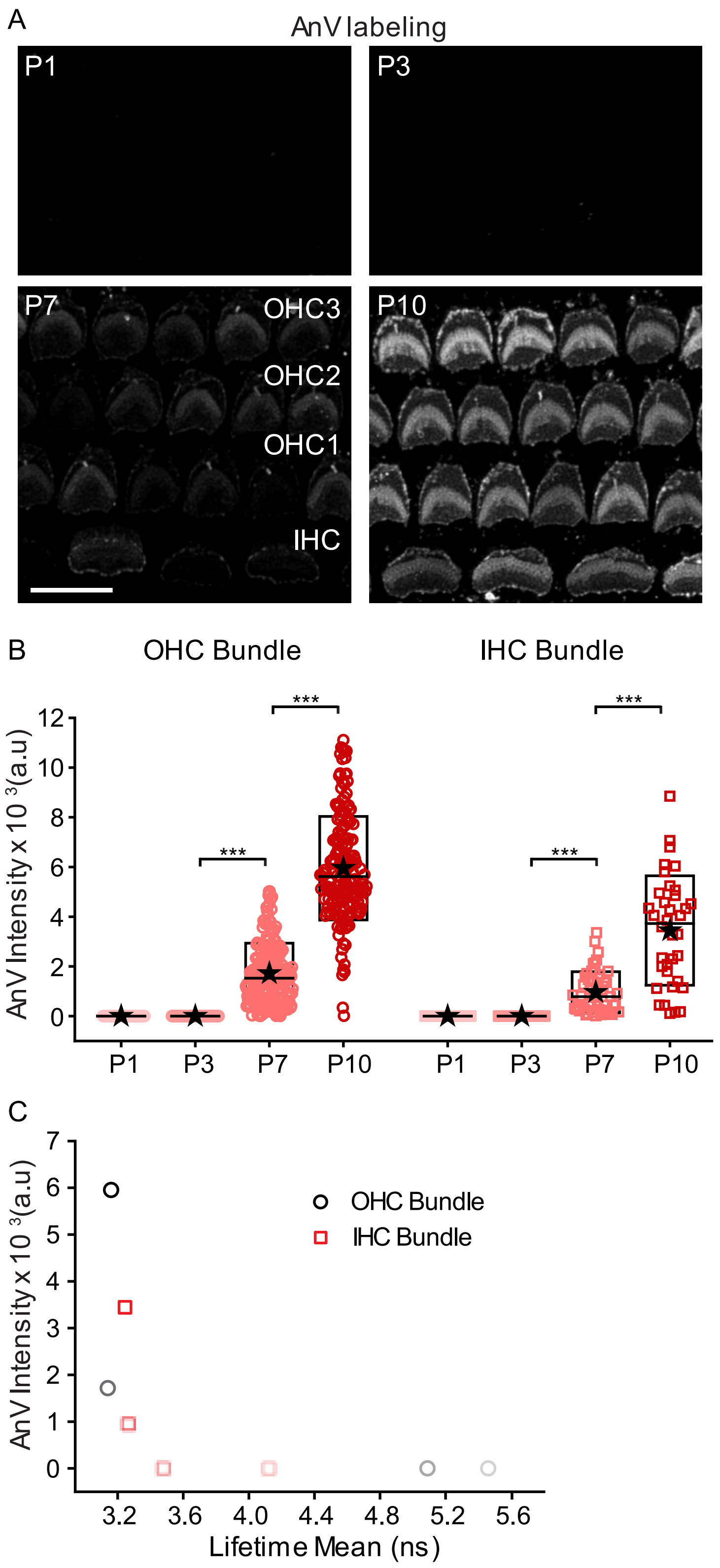


**Supplementary Figure 5:** A) Confocal images of OHCs and IHCs labelled with AnV from mid-apical turns of P1, P3, P7 and P10 rats. B) Quantification of AnV intensity for OHC and IHC bundles at different ages.

C) Graph obtained by plotting AnV intensity on the y-axis and the lifetime measured on the x-axis for OHB (circles) and IHB (squares) at different ages with darker shade indicating older ages. Scale bar = 10 μm.

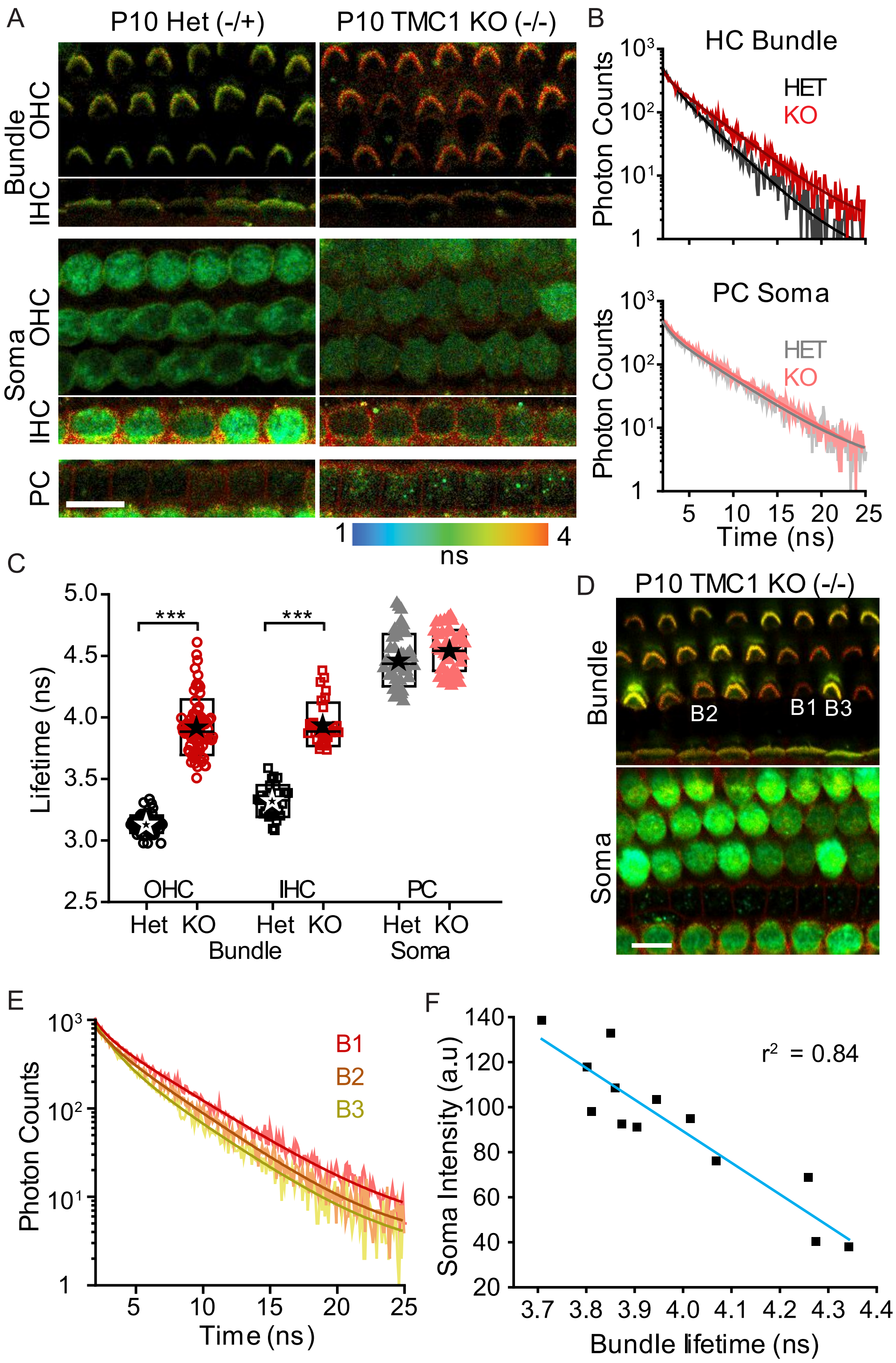


**Supplementary Figure 6:** A) FLIM images from mid-apical turn of organ of Corti from P10 heterozygous and TMC1 KO mice focusing on the hair bundles of OHCs and IHCs (top 2 rows) and soma of OHCs, IHCs and PCs (rows 3, 4 and 5). B) Fluorescence decay data and the corresponding fitting curves comparing the HC bundles (top) and PC soma (bottom) from KOs (black and gray) and hets (red and salmon). C) Summary box plot showing the measured lifetime for the OHC and IHC bundles and PCs from the KOs and the controls. D) An example FLIM image of the hair bundles (top) and the corresponding soma (bottom) from a P10 mid-apical turn showing the range of lifetimes and the soma intensity seen from the neighboring HCs. E) The fluorescence decay data and the corresponding fitting curves for three HC bundles B1, B2 and B3 highlighted in D). F) The HC bundle lifetime and the soma intensity of the corresponding HC was measured for the images shown in D) and plotted to show a strong correlation between both ($r^2 = 0.84$, $p < 0.001$).

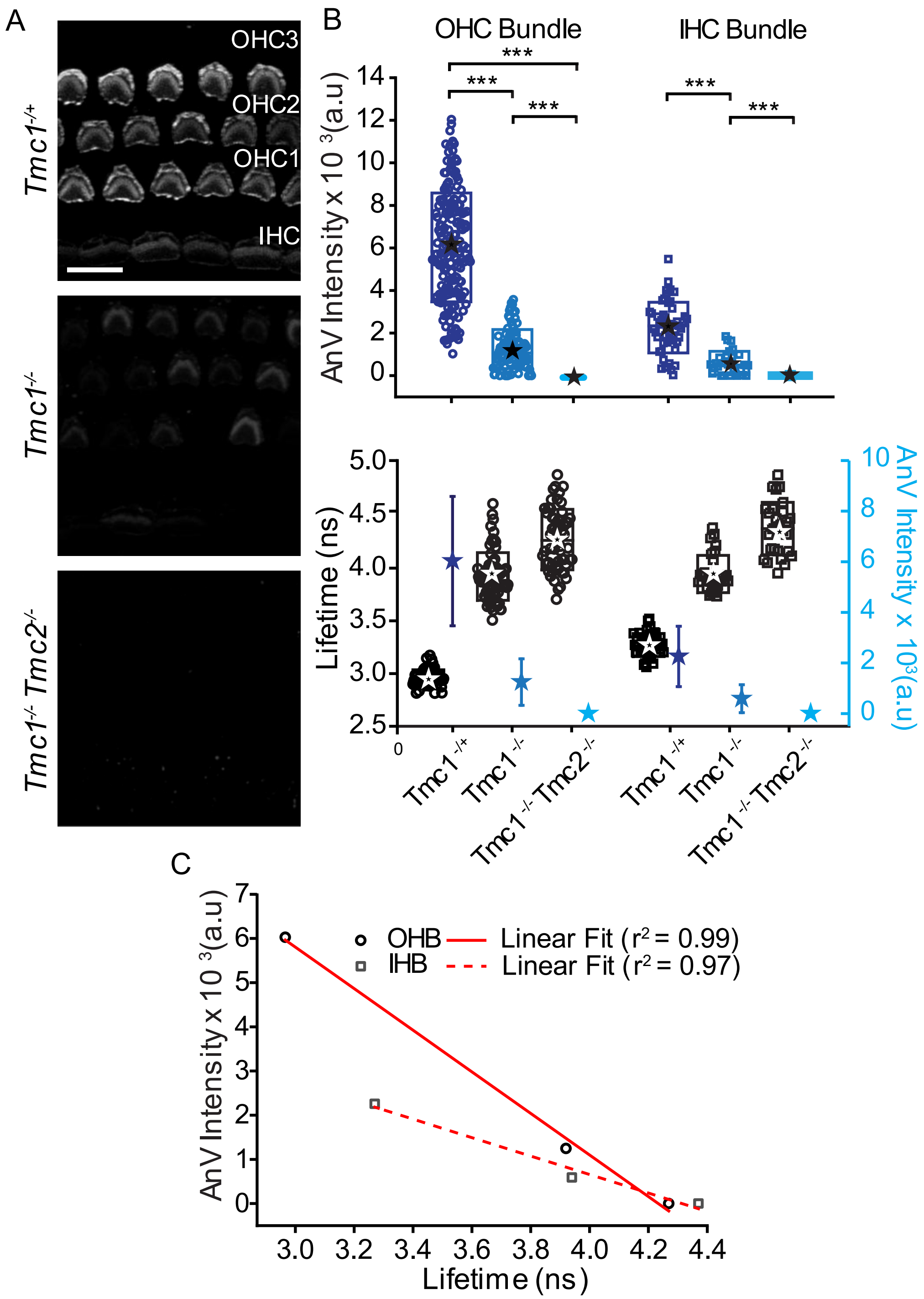


**Supplementary Figure 7:** A) Confocal images of OHCs and IHCs labelled with AnV from mid-apical turns of P10 *Tmc1*$^{-/+}$, *Tmc1*$^{-/-}$ and *Tmc1*$^{-/-}$*; Tmc2*$^{-/-}$ B) Quantification of AnV intensity (top panel) and AnV intensity and lifetime plotted on the y-axis (bottom panel) for OHC and IHC bundles for *Tmc1*$^{-/+}$, *Tmc1*$^{-/-}$ and *Tmc1*$^{-/-}$*; Tmc2*$^{-/-}$. C) Plot of mean AnV intensity vs. mean lifetime measured for OHB (circles) and IHB (squares) of P10 *Tmc1*$^{-/+}$, *Tmc1*$^{-/-}$ and *Tmc1*$^{-/-}$*; Tmc2*$^{-/-}$. Scale bar = 10 μm.

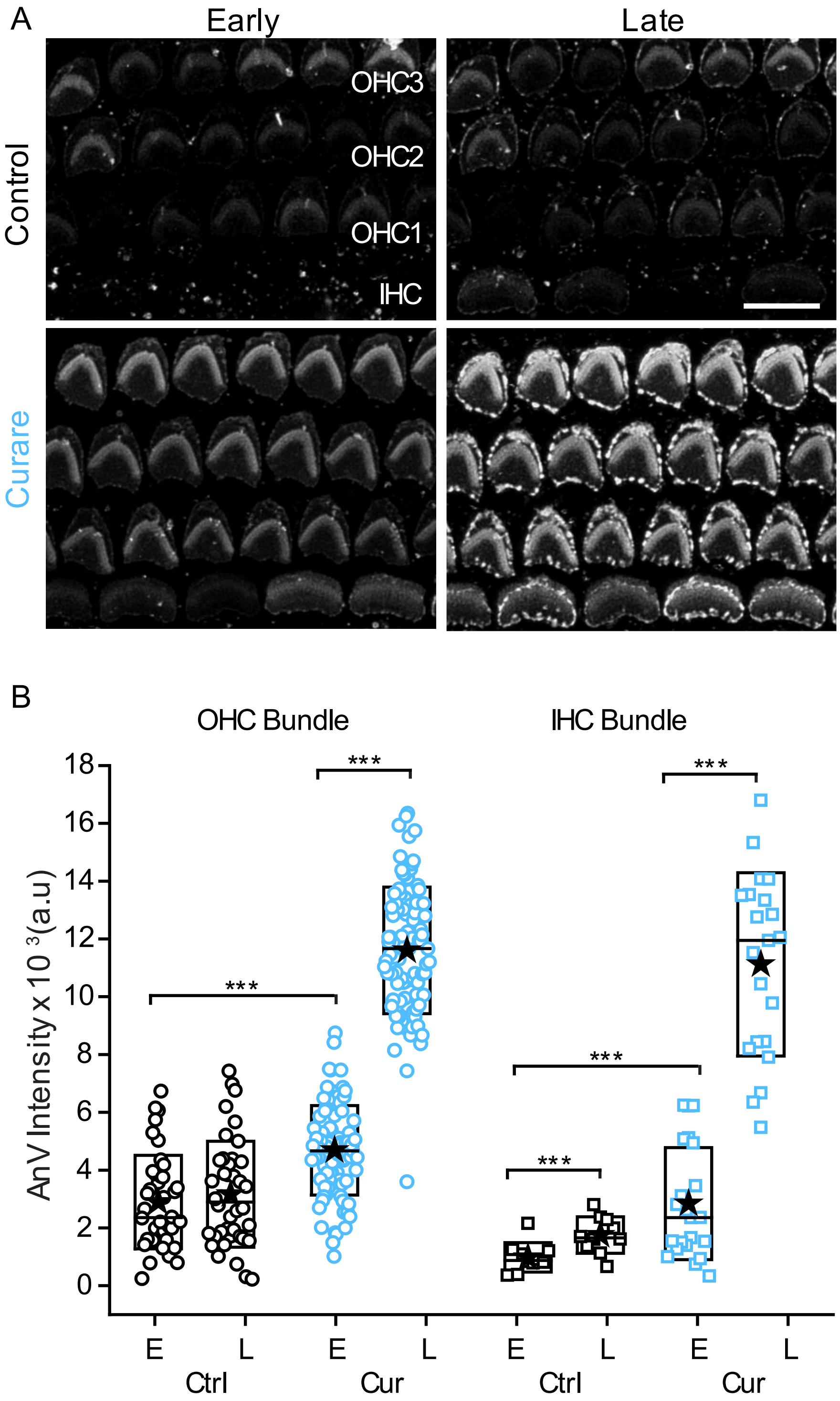


**Supplementary Figure 8:** A) Confocal images of OHCs and IHCs labelled with AnV from P10 rat mid-apical turns untreated (control) or treated with 1mM curare for 5-10 mins (early) or 30 mins (late) B) Quantification of AnV intensity for OHC and IHC bundles treated as in A. Scale bar = 10 $\mu$m.

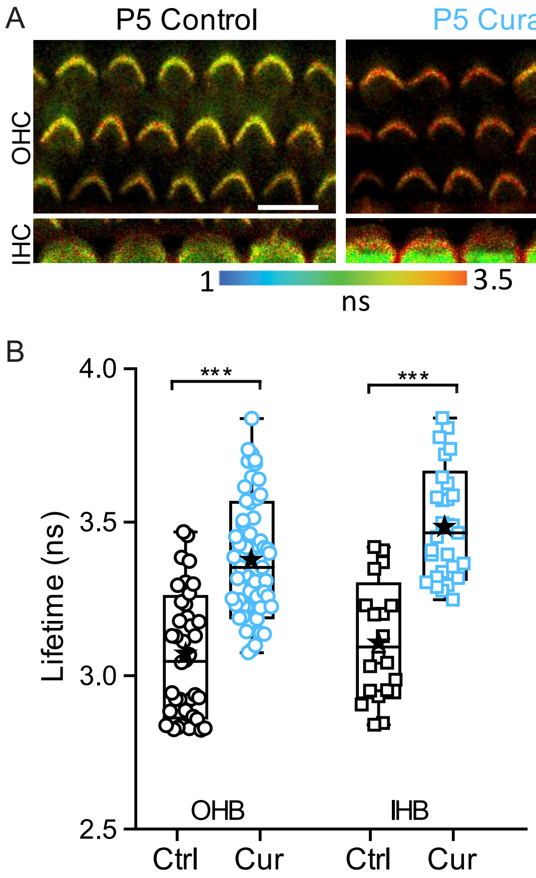


**Supplementary Figure 9:** A) FLIM images of OHCs and IHCs labelled with BODIPY 1c from P5 rat mid-apical turns untreated (control) or treated with 1mM curare for 5-10 mins (early) B) Quantification of lifetime for OHC and IHC bundles treated as in A. Scale bar = 10 μm.

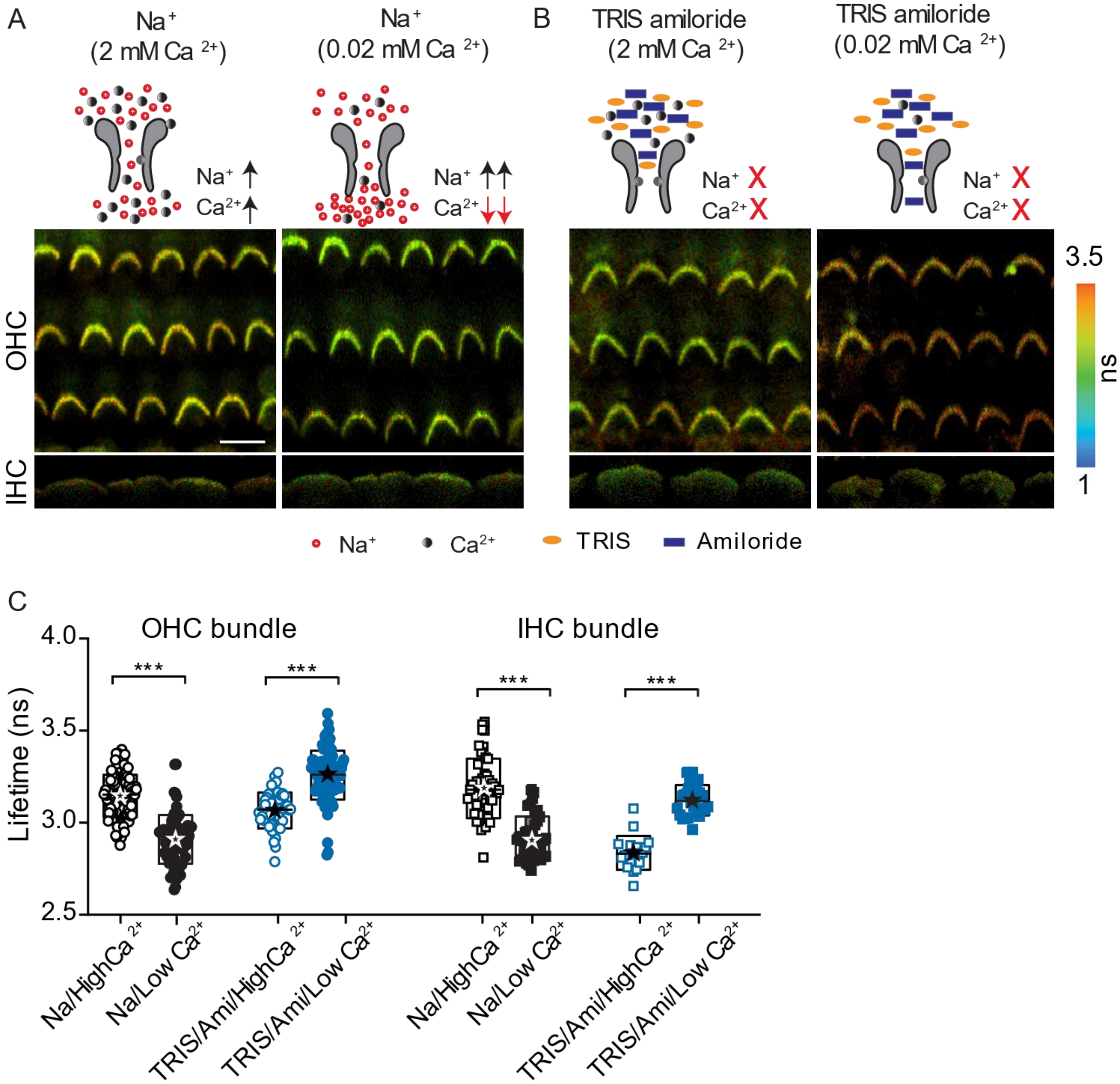


**Supplementary Figure 10:** A, B) FLIM images of OHCs and IHCs labelled with BODIPY 1c from P10 rat mid-apical turns in different external solution conditions, A) $Na^+$ with high (2mM) and low (0.02mM) $Ca^{2+}$ B) $TRIS^+$ and amiloride external with high and low $Ca^{2+}$ B) Quantification of lifetime for OHC and IHC bundles treated as in A and B. Scale bar = 10 μm.
